# Genomic and metabolic adaptations of biofilms to ecological windows of opportunities in glacier-fed streams

**DOI:** 10.1101/2021.10.07.463499

**Authors:** Susheel Bhanu Busi, Massimo Bourquin, Stilianos Fodelianakis, Grégoire Michoud, Tyler J. Kohler, Hannes Peter, Paraskevi Pramateftaki, Michail Styllas, Matteo Tolosano, Vincent De Staercke, Martina Schön, Laura de Nies, Ramona Marasco, Daniele Daffonchio, Leïla Ezzat, Paul Wilmes, Tom J. Battin

## Abstract

Microorganisms dominate life in cryospheric ecosystems. In glacier-fed streams (GFSs), ecological windows of opportunities allow complex microbial biofilms to develop and transiently form the basis of the food web, thereby controlling key ecosystem processes. Here, using high-resolution metagenomics, we unravel strategies that allow biofilms to seize this opportunity in an ecosystem otherwise characterized by harsh environmental conditions. We found a diverse microbiome spanning the entire tree of life and including a rich virome. Various and co-existing energy acquisition pathways point to diverse niches and the simultaneous exploitation of available resources, likely fostering the establishment of complex biofilms in GFSs during windows of opportunity. The wide occurrence of rhodopsins across metagenome-assembled genomes (MAGs), besides chlorophyll, highlights the role of solar energy capture in these biofilms. Concomitantly, internal carbon and nutrient cycling between photoautotrophs and heterotrophs may help overcome constraints imposed by the high oligotrophy in GFSs. MAGs also revealed mechanisms potentially protecting bacteria against low temperatures and high UV-radiation. The selective pressure of the GFS environment is further highlighted by the phylogenomic analysis, differentiating the representatives of the genus *Polaromonas*, an important component of the GFS microbiome, from those found in other ecosystems. Our findings reveal key genomic underpinnings of adaptive traits that contribute to the success of complex biofilms to exploit environmental opportunities in GFSs, now rapidly changing owing to global warming.

## Introduction

Ecosystems and their constituent biota are finely tuned to the seasonal variations of their environment. This phenology is particularly pronounced in glacier-fed streams (hereafter GFSs), which are commonly enveloped by snow cover and darkness in winter, and subject to high flow and sediment mobilization in summer. Yet, ecological ‘windows of opportunity’ arise in spring and autumn^1,2^, and are characterized by elevated light and nutrient (N, P) availability along with moderate flow, allowing algae and cyanobacteria to rapidly develop ‘green oases’ of phototrophic biofilms. Partially due to the absence of terrestrial organic matter subsidies from the catchment, this punctuated exploitation of solar energy in an otherwise energy-limited ecosystem, transiently forms the base of the food web and ecosystem energetics^1,3^. Such windows of opportunity may therefore function as ‘ecosystem control points’^4^ with disproportionately high ecological processing rates affecting ecosystem dynamics relative to longer intervening time periods. These ecosystem control points are widely distributed across ecosystems and vary across spatial and temporal scales^4^. However, our understanding on the microbiology of the communities that facilitate ecosystem control points remains limited to date.

Owing to climate change, the mass balance and melting dynamics of mountain glaciers are rapidly changing worldwide, altering the annual distribution of runoff in GFSs^5^. Invigorated glacial melt increases discharge and sediment delivery, but after glaciers shrink past a certain point (*i.e*., ‘peak water’), GFSs are likely to become warmer, less turbid, and less hydrologically dynamic^6^. These changes are almost certain to have substantial impacts on GFS ecosystem structure and function by either contracting or extending the duration of these windows of opportunity. It is therefore critical to understand how benthic biofilms operate during these times in order to predict how these ecosystems are likely to change and operate in the future^6^.

In streams, and GFSs in particular, biofilms closely interact with the sedimentary environment. For instance, the extracellular polymeric substances (EPS) produced by biofilms can stabilize fine sediments^7^. On the other hand, larger sediments (*e.g.*, boulders) resist flow-induced disturbance, thereby conferring stability to biofilms^8^. This appears particularly important in GFSs characterized by notoriously unstable fine sediments. Furthermore, light is minimally attenuated within a thin layer of water flowing over protruding boulders, which facilitates photosynthesis and biofilm growth. Therefore, it is particularly advantageous for phototrophic biofilms to colonize boulders, which form islands of stability in an otherwise highly unstable ecosystem. This may allow them to persist even during unfavorable periods, and potentially provide a model for microbial life during the windows of opportunity.

The relationship between photoautotrophs, such as algae and cyanobacteria, and microorganisms, primarily other bacteria, regulates nutrient and carbon cycling, and therefore represents a fundamental ecological interface in aquatic ecosystems. This interface (*i.e.*, the phycosphere) has received substantial attention in pelagic ecosystems over the last decades^9–12^, but less so in stream ecosystems. While early work on phototrophic biofilms colonizing the benthic zone in streams has highlighted the role of algal-bacterial interactions for carbon and nutrient fluxes^13,14^, we do not currently understand the fine-scale mechanisms of such interactions. For example, cyanobacteria produce pigments that protect biofilms against harmful UV-radiation^15^, while mucilage-rich algal colonies (*e.g.*, *Hydrurus* spp.) provide labile organic matter to heterotrophic microorganisms and facilitate their attachment. Such interactions may foster facultative interactions between photoautotrophs and other microorganisms, which, similar to the phycosphere, may be particularly beneficial to microbial life in oligotrophic and harsh ecosystems such as GFSs. Unraveling the genomic and metabolic underpinnings of algal-bacterial relationships in biofilms helps to better understand a most successful mode of microbial life in an extreme ecosystem.

Here we dissect the microbiome of GFSs and describe unprecedented genomic underpinnings of the adaptive mechanisms that contribute to the success of complex biofilms. Using 16S rRNA and 18S rRNA genes amplicon sequencing, we assess the microbiome structure of biofilms associated with two sedimentary habitats that are common in GFSs, namely sandy sediments (*i.e.*, epipsammic biofilms) and boulders (*i.e.*, epilithic biofilms). Furthermore, using high, genome-resolved metagenomics, we screen twenty-one epilithic biofilm microbiomes for energy pathways and cross-domain metabolic interactions. Our findings suggest the diversification of energy-acquiring pathways and metabolic interactions as relevant for epilithic biofilms to adapt to these ecological windows of opportunity, which are likely to become more prevalent as glaciers worldwide recede.

## Results and Discussion

### Sedimentary habitats affect microbiome structure and assembly

We used 16S rRNA and 18S rRNA gene amplicon sequencing to compare the microbiome structure of 48 epipsammic and epilithic biofilm samples from GFSs in the New Zealand Southern Alps (NZ) and in Caucasus (CC) (*Methods*) (Fig. 1a; Supp. Fig. 1a-b). These geographically distant streams, transcending hemispheres, were selected in order to draw more generalisable conclusions about microbiome structure and assembly. Moreover, to have comparable samples, the collection was largely constrained to the vernal and autumnal windows of opportunity. We found that both the prokaryotic and eukaryotic communities differed between the two habitat types in terms of community structure and alpha diversity (Fig. 1b-c). Overall, taxonomic differences were even apparent at the phylum level, despite high inter-sample variability within the categories (Supp. Fig. 1c-d). Geography explained 11.5% and 12.9% of the variability in the prokaryotic and eukaryotic datasets (db-RDA, *p* < 0.05 for both datasets), while sedimentary habitats explained an additional 10% and 8.3% of the variability (db-RDA, *p* < 0.05 for prokaryotes and eukaryotes).

**Figure 1.**
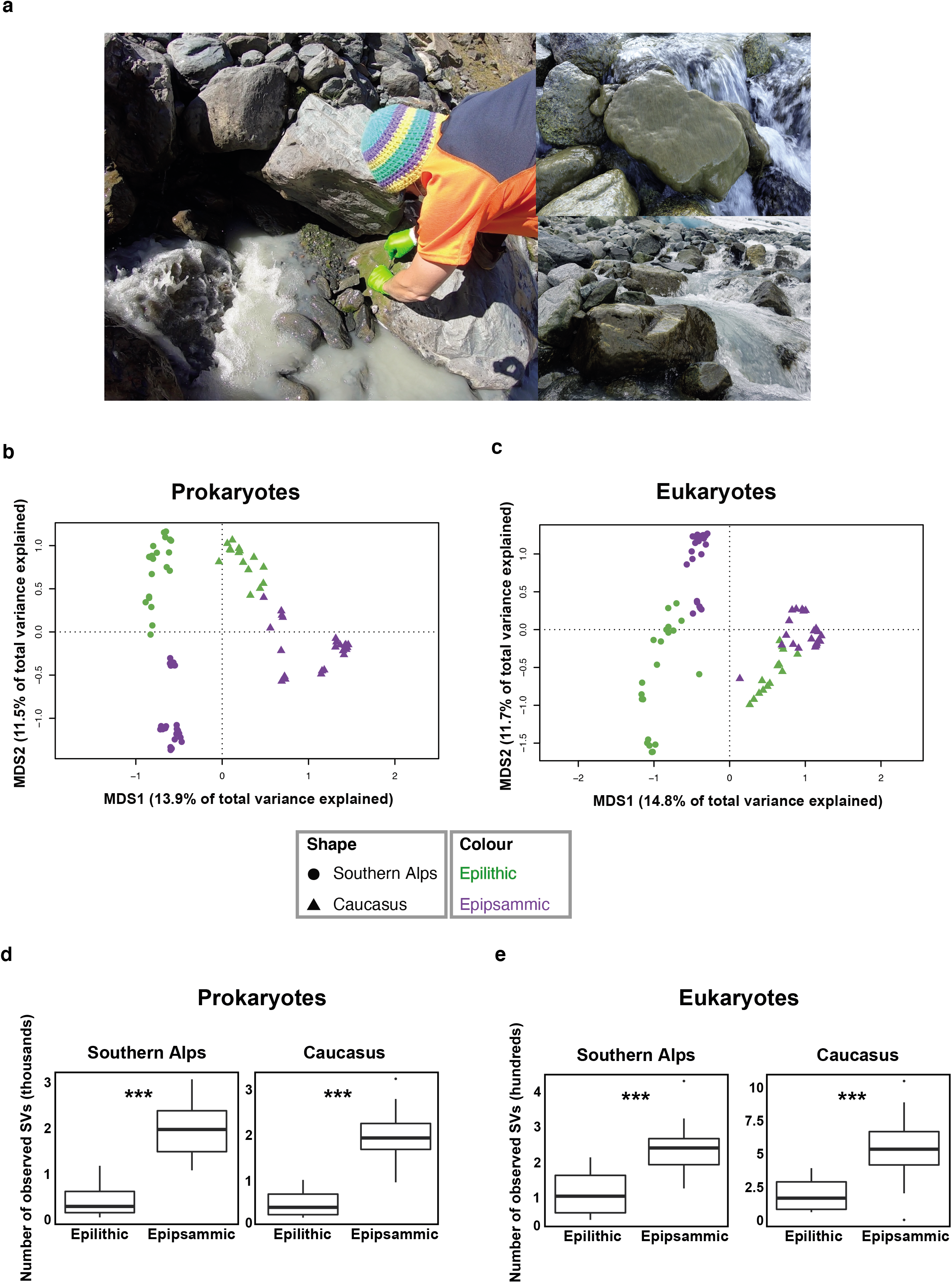
Sedimentary habitats affect microbiome structure and assembly. (a) Representative images of sample collection indicating GFS and adjacent epilithic biofilm (left) with images of epilithic biofilms (right). Photo credits: Martina Schön. Ordination analyses of the epipsammic and epilithic biofilm based on prokaryote (b) and eukaryote (c) metabarcoding profiles from Southern Alps and Caucasus. Microbial richness across geographic locations and sample types in (e) prokaryotes and (f) eukaryotes.

The estimated α-diversity (*i.e.*, species richness of amplicon sequence variants; ASVs) was higher for both prokaryotes and eukaryotes in epipsammic biofilms when compared to epilithic biofilms (2-3 fold differences, non-parametric t-tests, *p* < 0.001) (Fig. 1d-e). It is plausible that dispersal facilitated by the transport of fine sediments from various upstream sources (*e.g.*, subglacial environment, bare rock and soils) leads to the greater diversity of the epipsammic biofilms. Overall, our results unravel distinct microbiome structures for both habitats within the same GFS reaches; our results thereby agree with previous studies demonstrating the relationship between streambed physical variation and spatial biodiversity dynamics^16,17^. Streambeds, including their biofilms, are understood as landscapes where dispersal among patches can shape biodiversity and resilience^18–20^. Therefore, we hypothesized that epilithic communities are partially structured by dispersal from epipsammic communities that typically dominate the GFS streambeds by area. Using Sloan’s neutral community model^21^, we instead found that the composition of the epilithic biofilms is not dictated by a source-sink relationship with the epipsammic communities (*Supplementary text*). In other words, the epilithic biofilm communities are not determined by epipsammic communities that typically surround the boulders within the complex landscape of the GFS streambed.

### Metagenomics unveils the complexity of epilithic biofilms

To unveil the full complexity of the epilithic biofilms, we performed whole genome shotgun metagenomics on 21 epilithic samples from four GFSs each in NZ and CC (Supp. Fig. 1a-b); low biomass associated with sandy sediments precluded epipsammic biofilms from metagenomic analysis. High-resolution sequencing, after quality filtering yielded on average 1.2 × 10^8^ (± 1.4 ×10^7^ s.d.) reads per sample which were assembled to obtain an average of 8.7 × 10^5^ contigs per sample, that were subsequently binned. Bacteria and eukaryotes dominated the biofilm communities across all samples (Supp. Fig. 2a). Seventy-three (70 bacteria and 3 archaea) medium-to-high quality (>70% completion, <5% contamination) metagenome-assembled genomes (MAGs) from a total of 662 MAGs formed the pool of the prokaryotes. As seen from the phylogenomic analysis, the high-quality MAGs span the bacterial tree of life and based on this, along with the taxonomic information, many of these plausibly represent novel species (Fig. 2a). Aggregated at the genus level, *Polaromonas* was both abundant and prevalent in the biofilms along with representatives of *Flavobacterium*, *Cyanobacteria*, and unclassified MAGs from the Bacteroidota and Candidate Phyla Radiation (CPR; *Patescibacteria*) (Fig. 2b). These taxa were found in over half of the samples, irrespective of geographic origin. The CPR bacteria have only recently been identified based on genomic data^22^, and *Patescibacteria* specifically have been reported from oligotrophic ecosystems, including groundwater^23^ and thermokarst lakes^24^. Their apparently minimal biosynthetic and metabolic pathways may help them dwell in these ecosystems, which is of equal relevance in GFSs.

**Figure 2.**
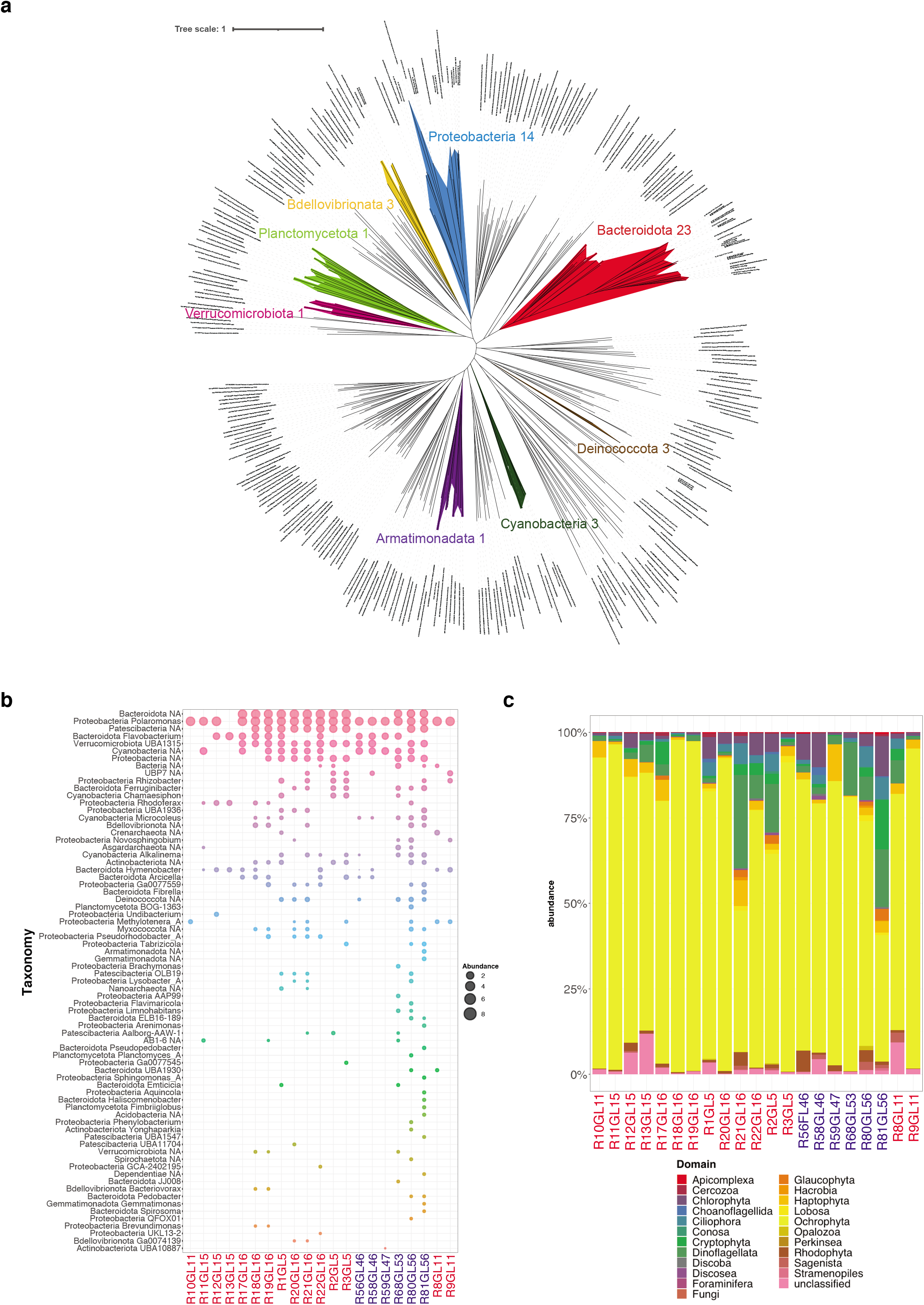
Metagenomics unveils the complexity of epilithic biofilms. (a) Bacterial phylogenetic tree constructed using high-quality (>90% completion and <2% contamination) MAGs reconstructed from the epilithic biofilms. The numbers beside the phylum names indicate the number of high-quality MAGs assigned to the respective phylum. (b) Normalized abundance of reconstructed prokaryotic genomes, *i.e.*, MAGs, from the epilithic biofilms. Taxonomy at phylum and genus levels is depicted. NA: unclassified genus. Samples from the Southern Alps are indicated in red, while those from Caucasus are shown in blue. (c) Eukaryotic relative abundance profile obtained from metagenomic sequencing across all epilithic biofilms samples.

Alongside these bacteria, archaea contributed less than 1% to the microbiome of epilithic biofilms, with representatives of Asgardarchaeota, Crenarchaeota and Nanoarchaeota. Intriguingly, the recently discovered lineages of Asgardarchaeota^25,26^ have been reported from freshwater sediments, yet not from cryospheric environments. Algae, mostly diatoms and *Hydrurus* (Ochrophyta phylum), as well as dinoflagellata were the most important photoautotrophs of the eukaryotic domain (Fig. 2c). The prevalence of *Hydrurus* (~87% relative abundance) underscores the function of this filamentous alga as a resource to higher trophic levels in GFS^27^. Our metagenomic insights further support the notion that phototrophic biofilms are highly diverse with representatives from all three domains of life^28^.

In addition to the archaeal, bacterial and eukaryotic community members, we also found a diverse viral community associated with epilithic biofilms (Supp. Fig. 2b). Most of the viruses were bacteriophages targeting abundant MAGs such as *Flavobacterium*, *Pseudomonas*, and *Bacillus* genera, but we also identified eukaryotic phages (*i.e*., *Paramecium bursaria* Chlorella virus). Few have studied viruses in stream biofilms to date^29^, potentially because it was common wisdom that the biofilm mode of life protects bacteria from viral infection. While viruses have previously been shown to be abundant in glaciers^30,31^, our findings are the first to provide evidence for a diverse and likely active viral community in GFS biofilms where they may influence bacterial growth and both carbon and nutrient cycling as on the glacier surface^30^.

### Epilithic biofilms form the basis for a ‘green’ food web

Cyanobacteria and eukaryotic algae figured among the most important photoautotrophs in the epilithic biofilms that form the basis of the ‘green’ food web during the window of opportunity. While these photoautotrophs are well known to use chlorophyll to capture solar energy, little is known on retinal-based phototrophy using rhodopsins in GFSs. Intriguingly, we found that MAGs from sixteen out of twenty phyla in the epilithic biofilms, including the abundant groups, such as Proteobacteria (*Polaromonas*) and Bacteroidota (*Flavobacterium*), encoded for (bacterio-)rhodopsins (Fig. 3a). These also included genes encoding for light-harvesting complex 1 (LH1), reaction centre (RC) subunits (*pufBALM*), and transcriptional regulators (*ppsR*) required for aerobic anoxygenic phototrophs along with rhodopsins as a signature of energy-limitation adaptations (Fig. 3a). Recently, rhodopsins were also reported to serve as a photoprotectant in *Flavobacterium* from glaciers^32^. Collectively, our findings unveil multiple strategies of photoautotrophy, which may help cyanobacteria and algae in maximising the exploitation of solar energy and to thrive during windows of opportunity.

**Figure 3.**
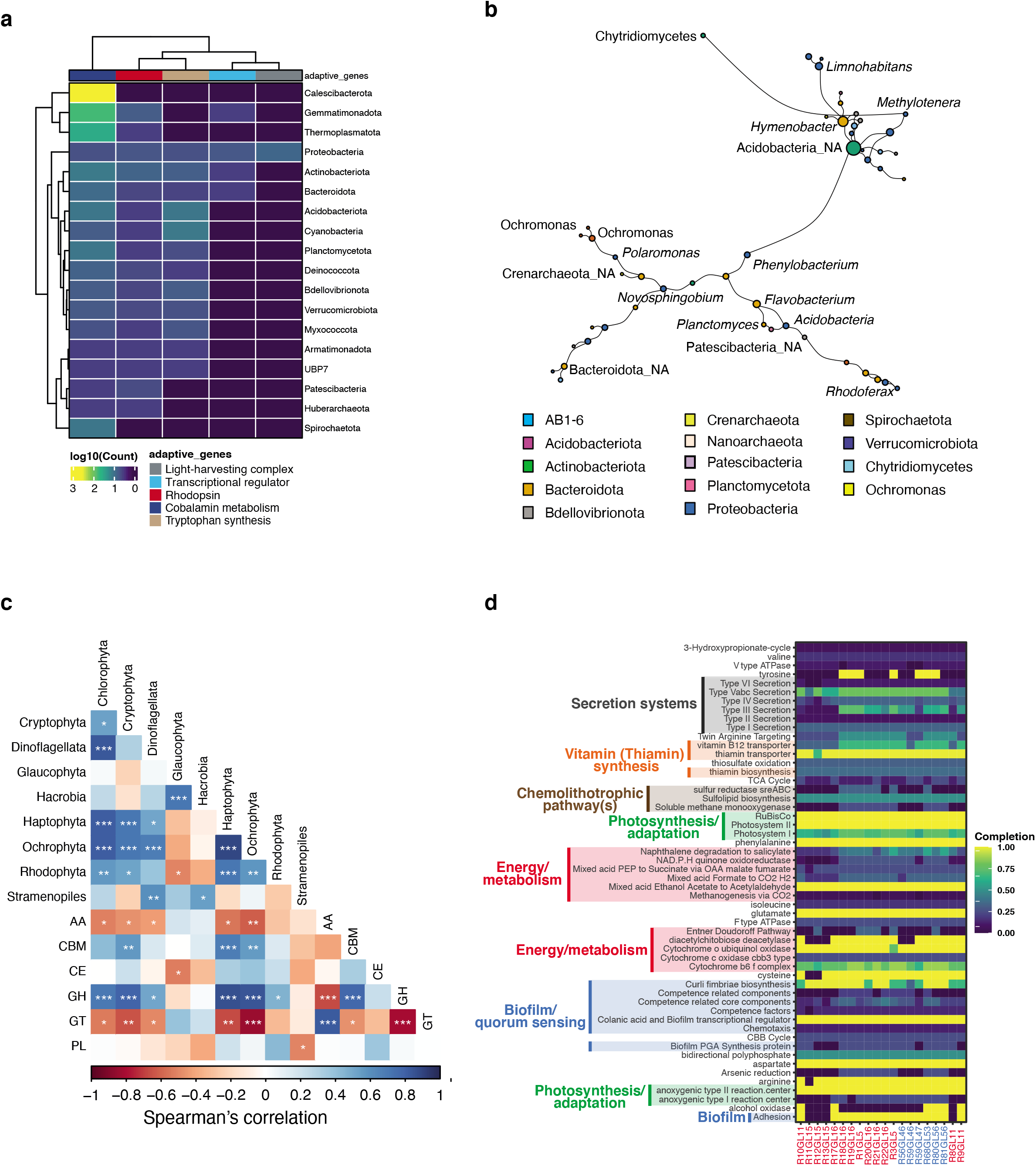
Epilithic biofilms are the basis for a ‘green food chain’. (a) Abundance of genes involved in energy production (*light-harvesting complex*, *transcriptional regulator for phototrophy*, and *rhodopsin*) and photo-heterotrophic interactions (*cobalamin metabolism* and *tryptophan synthesis*), across all prokaryotic phyla are represented in the heatmap. Values indicate the log10 abundance per gene within the phyla. (b) Largest component of the co-occurrence network between pro- and eukaryotic MAGs. Each node corresponds to a MAG (pro- or eukaryote). Size of the node corresponds to degree centrality and the edges represent the positive coefficients of correlation between each node. Colour of each node represents the phylum annotation. NA: unclassified genus. (c) Spearman’s correlation analyses of relative abundances of eukaryotic primary producers with the CAZyme abundances. FDR-adjusted *p*-values are indicated by *, *i.e.*, * < 0.05, ** < 0.01, *** < 0.001. (d) KEGG orthology (KO) pathways enriched in epilithic biofilms compared to publicly available cryospheric metagenomes were further assessed via KEGGDecoder for pathway completion and are displayed. The completeness of the pathways is indicated in the heatmap, per sample.

Rapid growth should be advantageous for primary producers, such as cyanobacteria, to best exploit the short windows of opportunity in GFSs. Moreover, functional independence from other microorganisms could thereby allow them to efficiently react to a window of opportunity. To test this hypothesis, we assessed the relationship between projected times of growth (doubling time in hours), with the median KEGG pathway completion within each MAG. Strikingly, most cyanobacterial MAGs (n = 38/44, 86%) exhibited decreased projected times of growth with respect to median KEGG module completion (Spearman’s correlation: r = −0.41, adj. *p* < 0.05). These observations suggest that when encoding all genes to form a complete KEGG pathway, phototrophic taxa within these epilithic biofilms may indeed be self-sufficient, thereby reducing their dependency from other (micro)organisms and fostering growth.

Given the energetic constraints in GFSs, it would be beneficial for bacterial heterotrophs to interact with these photoautotroph (micro)organisms for meeting their energy and nutrient demands. To investigate such cross-domain relationships, we used network analyses and identified key interacting taxa based on positively co-occurring nodes, using all prokaryotic and eukaryotic MAGs (see *Methods*). Based on a null model assessment (see *Methods*), our interaction networks showed preferential attachment within the nodes, along with increased centralities (*i.e.*, degree and edge-betweenness, Supp. Fig. 3a-b), suggesting that the interactions within these networks were not random. More importantly, the largest connected component (based on degree and betweenness centralities) of the interaction network contained taxa spanning archaea, bacteria and eukaryotic domains (Fig. 3b and Supp. Fig. 3b). Though *Acidobacteria* had a high degree of centrality, both *Polaromonas* and *Methylotenera* demonstrated strong interactions (> 0.6 betweenness centrality) with primary producers (including eukaryotic algae) and fungi. Specifically, *Polaromonas* had a strong interaction with algae, while *Methylotenera* co-occurred with *Chytridiomycetes* (Fig. 3b). These results support our hypothesis of heterotrophic bacteria co-occurring with eukaryotes, primarily algae, for metabolic cross-feeding, similar to those occurring in the phycosphere^10^.

Furthermore, our results hint at the existence of a more cryptic interaction in epilithic biofilms between parasitic fungi *Chytridiomycetes* and algae (*i.e*., *Ochrophyta*). Fungal parasitism on pelagic algae has been recently reported to be more important than expected, even with consequences for carbon and nutrient cycling as mediated by the fungal shunt^33,34^. The possibility of fungal parasitism on algae in epilithic biofilms further underlines the role of photoautotrophs as the foundation of a complex food web in GFSs as a typically energy-limited ecosystem.

### Genomic underpinnings of algae-bacteria metabolic interactions

As photoautotrophs grow and senesce, they increasingly exude intracellular material into their ambient environment, where it can be metabolized by heterotrophic bacteria through the action of extracellular enzyme activity (EEA). To explore this metabolic cross-feeding between bacterial heterotrophs and algae, we assessed the MAGs for genes encoding five common EEAs required for cleaving complex polysaccharides, phosphomonoesters and proteins^35^. Not unexpectedly, these genes were predominantly associated with bacterial heterotrophs, rather than with the photoautotrophs (Supp. Fig. 4), which suggests adapted genomic traits to meet specific metabolic needs of the heterotrophs. However, based on the presence of the EEA genes, especially among Cyanobacteria, we cannot discount the possibility of mixotrophy in the epilithic biofilms (Supp. Fig. 4a), including in other abundant members of the epilithic microbiome (Supp. Fig. 1c-d). The widespread occurrence of mixotrophy in planktonic communities, including Cyanobacteria, and the ensuing food web dichotomy is considered as an adaptive strategy to oligotrophic and cold ecosystems (*e.g*., the polar sea). Therefore, we argue that mixotrophy may also be an important trait of Cyanobacteria within GFS biofilms.

Carbohydrate-active enzymes (CAZymes) are the prime tools used by heterotrophic bacteria to initiate the degradation of polysaccharides, largely algae-derived in the GFS epilithic biofilms. To shed light on this potential trophic interaction identified through specific EEAs, we tested if all the CAZymes in the metagenomes covaried with the abundance of eukaryotes. Overall, we found positive correlations between eukaryote abundances and CAZymes, particularly carbohydrate-binding modules (CBM) and glycoside hydrolases (GH) (Supp. Fig. 4d). More specifically, these correlations were particularly pronounced for GH and some of the algal groups (*e.g*.,Ochrophyta, Haptophyta, Cryptophyta) that we found at relatively high abundances in the epilithic biofilms (Fig. 4c). As some of these algae are known to copiously produce sulfated carbohydrates^36^, we suggest a similar involvement of CAZymes (Supp. Table 1) in relation to polysaccharide degradation in GFS epilithic biofilms as recently reported from *Verrucomicrobia* isolates^37^. Given that sulfated carbohydrates are more resistant to bacterial degradation than other carbohydrates^37^, our findings suggest that they are still relevant to carbon turnover in an ecosystem that is inherently carbon limited.

**Figure 4.**
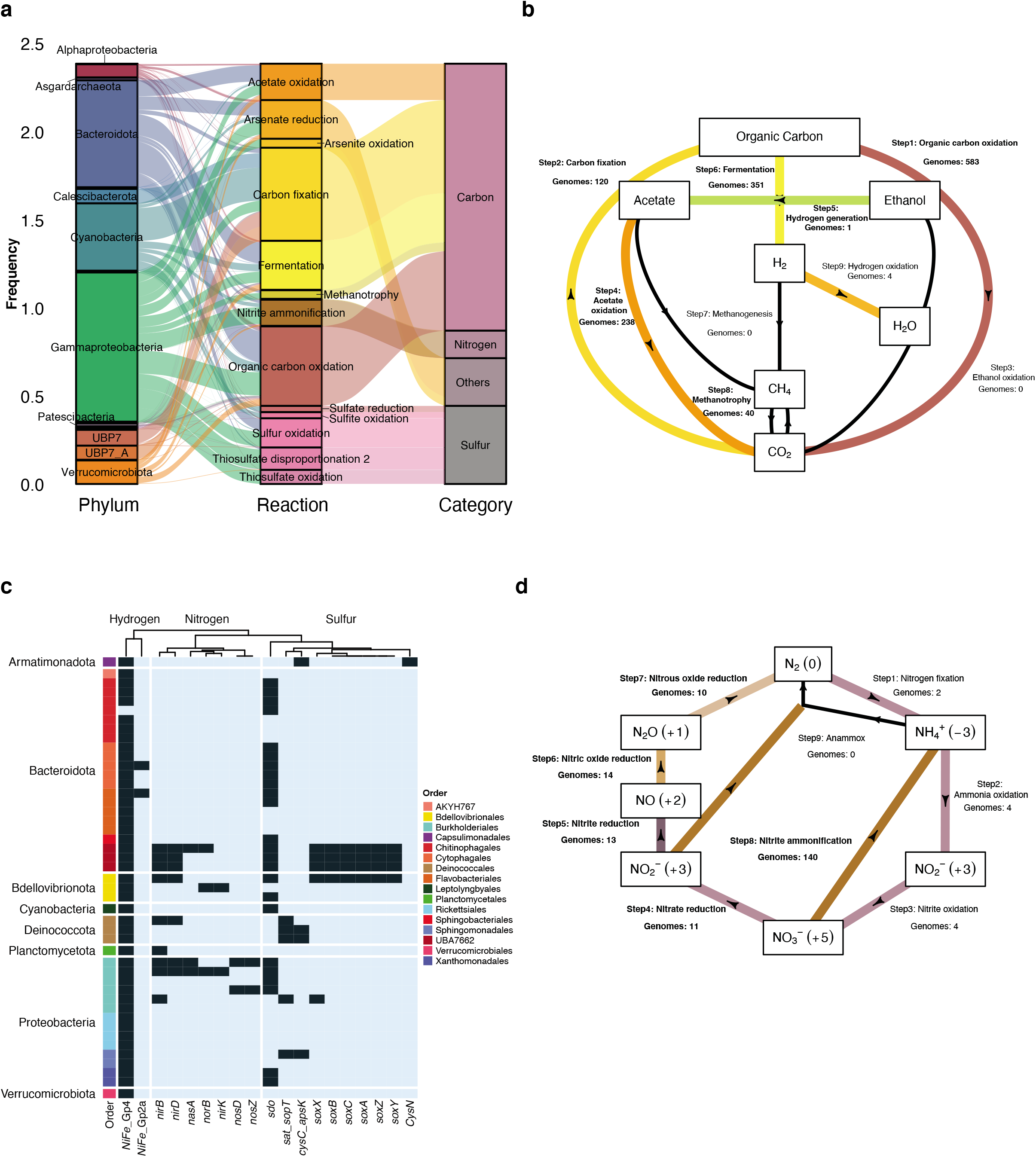
Functional redundancies across MAGs enable diverse energy acquisition and biogeochemical pathways. (a) The alluvial plot represents the metabolic pathways identified within all prokaryotic MAGs, with the respective taxonomic classification and category of nutrients. (b) Total number of MAGs encoding genes for and involved in the Carbon cycle (*see Methods*) are depicted in the flow gram created using a modified script from METABOLIC^91^. Each sub-pathway is indicated as a step with the corresponding number of genomes encoding the respective genes. (c) Phylum and order-level distributions of chemolithotrophic (hydrogen, nitrogen and sulfur) pathways with the respective gene copies per pathway are depicted in the heatmap. (d) Flow diagram indicating the MAGs encoding for pathways in the nitrogen cycle (*Methods*). Each sub-pathway is indicated as a step with the corresponding number of genomes encoding the respective genes.

In order to understand whether functions potentially geared towards cross-domain interactions were enriched solely in epilithic biofilms, we compared the KEGG orthology (KO) annotations from our metagenomes to 105 metagenomes from a wide range of ecosystems (Supp. Table 2). Strikingly, we found that KOs associated with quorum sensing, vitamin B12 (cobalamin) transporters and thiamine biosynthesis were enriched in epilithic biofilms compared to other ecosystems (Supp. Table 3). The associated pathways and their completion levels were evaluated using KEGGDecoder (Fig. 4d; Supp. Fig. 5) indicating a high completion of pathways associated with cross-domain interactions. These findings are in line with previous genomic insights into algal-bacterial interactions^38,39^, specifically with the observed upregulation of vitamin biosynthesis in bacteria (*Halomonas*) growing in the presence of algal extracts.

Furthermore, several MAGs were found to encode genes (*e.g*., quorum sensing, cobalamin metabolism, tryptophan synthesis) potentially facilitating algal-bacterial interactions (Fig. 4a). Particularly, cobalamin metabolism may be relevant for nutrient acquisition in algal-bacterial relationships^40^, whereas tryptophan was reported as a key signalling molecule involved in interactions between bacteria and associated phytoplankton^11,41^. Collectively these genomic insights stress cross-domain interactions as an adaptive potential that the epilithic microorganisms have developed to exploit the window of opportunity in GFSs.

### Energy acquisition and biogeochemical pathways in epilithic biofilm MAGs

The dominance (~88%) of MAGs encoding for organic carbon metabolism highlights the relevance of a ‘green food web’ during the windows of opportunity, potentially sustaining metabolic interactions between primary producers and heterotrophs. Further exploring the gene repertoire of the epilithic biofilms, we found that Cyanobacteria were one of the largest bacterial contributors to carbon fixation along with Bacteroidota and few Gammaproteobacteria (Fig. 4a). An in-depth analysis across the 662 MAGs revealed that 583 MAGs encoded genes involved in organic carbon oxidation, while 120 MAGs encoded genes involved in CO2 fixation. In line with the above findings, the majority of these MAGs was identified as Cyanobacteria along with few other phyla such as Proteobacteria, Asgardarchaeota, Crenarchaeota and Huberarchaeota. We also note that 351 MAGs encoded genes for fermentation (Fig. 4b) spanning several phyla, including Actinobacteriota, Bacteroidota, Patescibacteria, Planctomycetota and Verrucomicrobiota.

For biofilms to thrive in GFSs, even during the windows of opportunity, it appears opportune to diversify the exploitation of energy sources. Therefore, we performed an in-depth characterisation of chemolithotrophic pathways to disclose the potential role of minerals derived from the glacial comminution of bedrock as an energy source for microorganisms^42^. The prevalence of the *sox* gene cluster in representatives of the Bacteriodota (UBA7662) and Bdellovibrionota reveals the potential importance of inorganic sulfur oxidation in epilithic biofilms. This notion is supported by the broad occurrence of sulfur dioxygenases (SDOs) across the various phyla that facilitate sulfur oxidation (Fig. 4c). Interestingly, Tranter and Raiswell suggested that sulfates are derived from sulfide oxidation in comminuted bedrock^43^ potentially increasing sulfur availability and acquisition in glacial meltwaters^44^. Sulfide oxidation can stimulate carbonate weathering with the resulting CO2 potentially being fixed by algae and cyanobacteria in the epilithic biofilms — a link that appears relevant given that GFSs are often undersaturated in CO_2_^45^. Furthermore, we found that almost all MAGs encoded for group IV hydrogen dehydrogenases (NiFe_Gp4; Fig. 4c), which potentially serve as an alternate energy acquisition pathway. Hydrogen dehydrogenases have recently been reported to support primary production in various glacial and other extreme environments^46,47^. This suggests that lithogenic hydrogen may also contribute energy to bacteria within the epilithic biofilms.

Further genomic insights into the nitrogen cycle revealed the Dissimilatory Nitrate Reduction to Ammonium (DNRA, or nitrite ammonification) and, to a lesser extent, by denitrification, as major pathways (Fig. 4d). Relatively little is known regarding these two competing pathways in stream biofilms or sediments^48^. However, our insights into the cross-domain metabolic interactions suggest that epilithic algae provide significant amounts of organic carbon (*i.e*., electron donors), which may favour bacteria to grow using DNRA. This is in line with other ecosystems where DNRA is favoured over denitrification when alternate electron donors prevail over nitrate^49^. Our analyses revealed Burkholderiales (Gammaproteobacteria) as the largest contributor to nitrate assimilation and ammonia-oxidation genes (Fig. 5a). DNRA, if not conducive to N_2_O production, would enhance nitrogen recycling within epilithic biofilms through ammonia assimilation by algae and cyanobacteria, for instance. Our genomic evidence for nitrogen recycling that potentially overwhelms nitrogen losses through denitrification is corroborated by flux measurements from microbial mats in Antarctic GFSs^50^, and highlights recycling as a strategy to cope with nutrient limitation in glacier ecosystems^50–52^.

**Figure 5.**
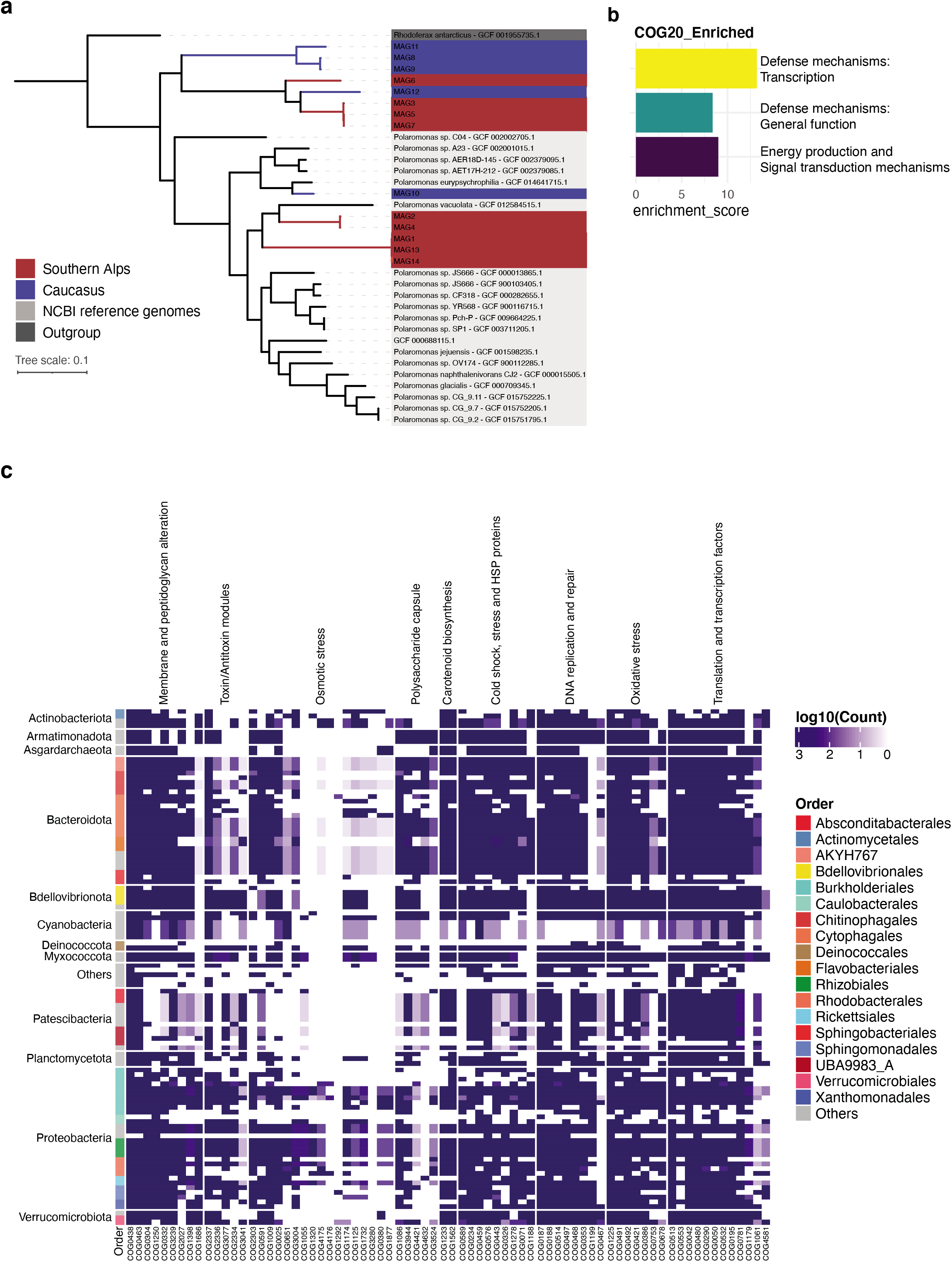
Genomic underpinnings of adaptation to the extreme GFS environment. (a) Phylogenomic tree based on *Polaromonas* genomes recovered from Southern Alps (red) and Caucasus (blue) along with publicly available genomes (grey) and an outgroup (*Rhodoferax*, dark grey). (b) Clusters of orthologous (COG20) group pathways enriched in epilithic biofilms MAGs compared to the reference genomes are depicted in the barplot. (c) Heatmap representing the abundance of genes involved in cold adaptation. Taxonomy at phylum and order levels is depicted. Columns indicate clusters of orthologous groups associated with adaptive genes.

Strikingly, we found only few MAGs, mostly belonging to Deinococcota, Gammaproteobacteria, Beijerinckiaceae and Crenarchaeota, involved in the oxidation of ammonia and nitrite potentially leading to the accumulation of nitrate. The involvement of archaea would be in line with recent studies showing ammonia oxidation by archaea in Arctic soils^53^ and with the observation that archaea couple ammonia oxidation with biomass formation (*i.e*., via CO_2_ fixation)^54^. Our finding that archaeal MAGs encode for carbon fixation genes (Fig. 4b) further highlight their role in ammonia oxidation and biomass accrual in epilithic biofilms. Overall, the overlap of metabolic capacities within the MAGs suggests that the epilithic biofilms may typify a ‘closed system’, where both carbon and nutrients are efficiently recycled.

### Genomic underpinnings of adaptation to the extreme GFS environment

The GFS environment is characterized by near-freezing temperatures, high UV-radiation, and high flow velocities. To assess potential adaptive traits of bacteria dwelling in epilithic biofilms, we first performed a phylogenomic analysis of *Polaromonas* spp., one of the most abundant and prevalent genera in the studied GFSs. Our analysis revealed that a few of the GFS *Polaromonas* formed clades that are distinct from *Polaromonas* identified in other environments (*Methods*), thus potentially comprising novel species’ (Fig. 5a). This phylogenomic pattern indicates that *Polaromonas* has evolved traits that facilitates its success in GFS, both in NZ and CC. To identify such traits, we created a pangenome and performed an enrichment analysis for clusters of orthologous genes. We found three categories that were significantly enriched in GFS *Polaromonas* compared to those from other environments (Supp. Table 4). Two categories are related to defense mechanisms, both general and transcription, and one to energy production (Fig. 5b). It is plausible that these mechanisms are related to high UV-radiation^55,56^ and oxidative stress^57^, as well as to cold stress responses as previously reported from other bacteria^58–60^. Furthermore, the presence of CRISPR-Cas proteins in the enriched clusters of orthologous genes (COGs) hint at defense mechanisms against phages (Supp. Table 4), which we showed to be present in the epilithic biofilms. This is in accordance with reports demonstrating that cryospheric bacteria (*Janthinobacterium* spp.) develop defense strategies, including biofilm formation^61^ and extracellular vesicle formation^62^ to escape viruses. On the other hand, the transcription of ‘defense mechanisms’ genes have been linked to cold adaptation in psychrophiles^58^. Cold-shock proteins regulate transcription at low temperature, while genes involved in membrane biogenesis^63^ and membrane transport proteins^64^, several of which are also enriched in the GFS *Polaromonas* genomes, are up-regulated. For example, in the psychrophilic *Colwellia psychrerythraea* 34H, adaptation to cold includes the maintenance of the cell membrane liquid-crystalline state via the expression of genes involved in polyunsaturated fatty acid synthesis^65^. Similarly, ATP-driven or proton motive secondary transport systems have been associated with solute transfers across membranes in bacteria and archaea as an adaptation to the cold^64^.

Our insights into the adaptive potential of *Polaromonas* to the GFS environment prompted us to expand our search for adaptive traits across all MAGs from the epilithic biofilms. Querying for 76 genetic traits spanning nine categories related to cold adaptation^59^, we found indeed distinct patterns of genomic adaptation across MAGs (Fig. 5c). Several MAGs encoded for genes associated with membrane and peptidoglycan alterations, cold and heat shock proteins, oxidative stress, and transcription/translation factors alongside DNA replication and repair. While all major phyla encoded for adaptive traits related to the outer membrane and cell wall, Proteobacteria were the predominant group with an overall higher copy number of genes involved in counteracting osmotic and oxidative stress. This is in line with metagenomic studies reporting an enrichment of sigma B genes in Antarctic mats, allowing for surviving severe osmotic stress during freezing^66^. Similarly, *Psychrobacter arcticus^67^* and *Planococcus halocryophilus* Or1^68^ were shown to have specific genomic modifications, particularly with genes involved in putrescine and spermidine accumulation, both of which are associated with alleviating oxidative stress. Furthermore, MAGs from Proteobacteria were characterized by high prevalence of genes potentially expressed in response to stressors, such as UV and reactive oxygen species (Fig. 5c).

In conclusion, our genome-resolved metagenomics analyses have set the stage for a mechanistic understanding of how the diversification of energy and matter acquisition pathways as well as metabolic interactions allow biofilms to thrive during windows of opportunity in GFSs. We acknowledge that a metagenomic time series outside and throughout windows of opportunity would be required to substantiate some of our observations. Nevertheless, our findings shed light on boulders as important habitats that confer stability of biofilms even outside the typical windows of opportunity. GFSs count among the ecosystems that are most vulnerable to climate change.

Therefore, our findings open a window into the future of how microbial life, with a strong photoautotrophic component, may look like in GFSs as glaciers shrink.

## Material and methods

### Sample collection

We sampled a total of eight GFSs from the New Zealand Southern Alps and the Russian Caucasus in early- and mid-2019, respectively, for a total of 27 epipsammic samples taken from sandy sediments and 21 epilithic biofilm samples from boulders adjacent to the epipsammic samples (Supp. Table 5). Epipsammic samples were collected from each GFS by first identifying three patches within a reach of ~5-10 m. From each patch, sandy sediments were taken from the <5 cm surface of the streambed with a flame-sterilized metal scoop and sieved to retain the 250 μm to 3.15 mm size fraction. While three epipsammic samples were taken from each stream, epilithic samples were taken opportunistically from up to three boulders per reach (Supp. Table 5). Epilithic biofilms were sampled using a sterilized metal spatula. All samples were immediately flash-frozen in liquid nitrogen in the field and transported and stored frozen pending DNA extraction. Streamwater turbidity, conductivity, temperature, and pH were measured *in situ* during the sampling (Supplementary Table 2).

### DNA extraction and purification

A previously established protocol^69^ was used to extract DNA from all samples. Briefly, 5 g of epipsammic and 0.05-0.1 g of epilithic biofilm were subjected to a phenol:chloroform-based extraction and purification method. The differential input volume for the DNA extractions were established to account for the differences in biomass between the epipsammic and epilithic biofilms. The samples were treated with a lysis buffer containing SDS along with 0.1 M Tris-HCl pH 7.5, 0.05 M EDTA pH 8, 1.25% SDS and RNase A (10 μl: 100 mg/ml). The samples were vortexed and incubated at 37 °C for 1 h. Proteinase K (100 μl; 20 mg/ml) was subsequently added and further incubated at 70 °C for 10 min. Samples were purified once with phenol/chloroform/isoamyl alcohol (ratio 25:24:1, pH 8) and the supernatant was subsequently extracted with a 24:1 ratio chloroform/isoamyl alcohol. Linear polyacrylamide (LPA) was used along with sodium acetate and ice-cold isopropanol for precipitating that DNA overnight at −20 °C. For epilithic biofilms, the entire protocol was adapted to a smaller scale due to the availability of higher DNA concentrations compared to sediment. The former was treated with 0.75 ml of lysis buffer (instead of 5 ml for sediment) and all subsequent volumes of reagents were adapted accordingly (see supplementary material). Furthermore, a mechanical lysis step of bead-beating was necessary along with a lysis buffer to facilitate DNA release from the more developed epilithic biofilms. Due to the higher DNA yields, the addition of LPA was omitted from the DNA precipitation step. DNA quantification was performed for all samples with the Qubit dsDNA HS kit (Invitrogen).

### Metabarcoding library preparation and sequencing

The prokaryotic 16S rRNA gene metabarcoding library preparation was performed as described in Fodelianakis *et al*.^70^, targeting the V3-V4 hypervariable region of the 16S rRNA gene with the 341F/785R primers and following Illumina guidelines for 16S metagenomic library preparation for the MiSeq system. The eukaryotic 18S rRNA gene metabarcoding library preparation was performed likewise but using the TAReuk454F-TAReukREV3 primers to target the 18S rRNA gene V4 loop^71^. Samples were sequenced using a 300-bp paired-end protocol partly in the Genomic Technologies Facility of the University of Lausanne (27 epipsammic samples) and partly at the Biological Core Lab of the King Abdullah University of Science and Technology (21 epilithic samples).

### Metabarcoding analyses

The 16S rRNA gene metabarcoding data were analysed using a combination of Trimmomatic^72^ and QIIME2^73^ as described in Fodelianakis *et al.^70^*, with the exception that here the latest SILVA database^74^ v138.1 was used for taxonomic classification of 16S rRNA and 18S rRNA gene amplicons. Non-bacterial ASVs including those affiliated to archaea, chloroplasts and mitochondria were discarded from the 16S rRNA amplicon dataset in all downstream analyses. ASVs observed only once were removed from both 16S rRNA and 18S rRNA amplicon datasets. Diversity analyses were performed in R using the *vegan^75^* and *metacoder^76^* packages. To test for a source-sink hypothesis from sediments to rocks, the Sloan’s Neutral Community Model^21^ was used based on the R implementation developed by Burns *et al*.^77^.

### Whole-genome shotgun libraries and sequencing

All epilithic biofilm DNA samples underwent random shotgun sequencing following library preparation using the NEBNext Ultra II FS library kit. Briefly, 50 ng of DNA was used for constructing metagenomic libraries under 6 PCR amplification cycles, following enzymatic fragmentation of the input DNA for 12.5 mins. The average insert size of the libraries was 450 bp. Qubit (Invitrogen) was used to quantify the libraries followed by quality assessment using the Bioanalyzer from Agilent. Sequencing was performed at the Functional Genomics Centre Zurich on a NovaSeq (Illumina) using a S4 flowcell.

### Metagenomic preprocessing, assembly, binning, and analyses

For processing metagenomic sequence data, we used the Integrated Meta-omic Pipeline (IMP)^78^ workflow to process paired forward and reverse reads using version 3.0 (commit# 9672c874; available at https://git-r3lab.uni.lu/IMP/imp3), as previously described^79^. IMP’s workflow includes pre-processing, assembly, genome reconstructions and additional functional analysis of genes based on custom databases in a reproducible manner. Briefly, adapter trimming is followed by an iterative assembly using MEGAHIT v1.2.9^80^. Concurrently, MetaBAT2 v2.12.1^81^ and MaxBin2 v2.2.7^82^ are used for binning in addition to an in-house method established previously^79^ for reconstructing metagenome-assembled genomes (MAGs). Binning was completed by selecting a non-redundant set of MAGs using DASTool^83^ based on a score threshold of 0.7. The quality of the MAGs was assessed using CheckM v1.1.3^84^, while taxonomy was assigned using the GTDB-toolkit v1.4.1^85^.

For the downstream analyses including identification of viruses, VIBRANT v1.2.1^86^ was used on the metagenomic assemblies. The output from this was used to identify the viral taxa using vConTACT2 v0.9.22^87^. Independently, the viral contigs were also validated using CheckV v0.7.0^88^. To estimate the overall abundances of eukaryotes along with prokaryotes including archaea, we used EUKulele v1.0.5^89^ with both the MMETSP and the PhyloDB databases, run separately, to confirm the detected eukaryotic profiles. To understand the overall metabolic and functional potential of the metagenome and reconstructed MAGs we used MANTIS^90^. Additionally, we used METABOLIC v4.0^91^, metabolisHMM v2.21^92^, and Lithogenie from MagicLamp v1.0 (https://github.com/Arkadiy-Garber/MagicLamp) to identify metabolic and biogeochemical pathways relevant for determining nutritional phenotypes of all MAGs along with the ‘*anvi-estimate-metabolism*’ function from anvi’o^93^. This information was manually validated based on the different tools to identify which MAGs encode for the respective pathways. Subsequently, to determine the growth rates of prokaryotes, we used codon usage statistics for detecting optimization of genes that are highly expressed, as an indicator of maximal growth rates with gRodon v1.0^94^. All the parameters, databases, and relevant code for the analyses described above are openly available at https://git-r3lab.uni.lu/susheel.busi/nomis_pipeline and included in the Code availability section.

### Eukaryote assembly and binning

To obtain eukaryotic MAGs, an alternate, custom pipeline (https://github.com/Mass23/NOMIS_ENSEMBLE/tree/coassembly) was established for coassembling the twenty-one epilithic biofilm sequence data with subsequent binning. Individual samples were first preprocessed similar to the workflow used in IMP, i.e., using FastP v0.20.0^95^. Subsequently, the reads were deduplicated to avoid overlap and enhance computation efficiency using *clumpify.sh* from the BBmap suite v38.79^96^. Thereafter, any reads mapping to bacteria or viruses were removed by filtering the reads against a Kraken2 v2.0.9beta^97^ maxikraken database available at https://lomanlab.github.io/mockcommunity/mc_databases.html. Only reads that were unknown or mapping to eukaryotes were retained and concatenated. This was followed by another round of deduplication using *clumpify.sh*. The concatenated reads were assembled using MEGAHIT v1.2.7 with the following options: *--kmin-1pass -m 0.9 --k-list 27,37,47,57,67,77,87 --min-contig-len 1000.* Following assembly, EukRep v0.6.7^98^ was used for retrieving eukaryotic contigs with a minimum length of 2000 bp and the ‘*-m strict*’ flag. These contigs were used for binning into MAGs as described herein.

Eukaryotic MAGs were binned using CONCOCT v1.1.0^99^. To do this, coverages were estimated for the contigs by mapping the reads of all samples against the contigs using the coverm v0.6.1 (https://github.com/wwood/CoverM) to generate bam files. These files were then used to generate a table with coverage depth information per sample. The protein coding genes of the MAGs was predicted with MetaEuk v4.a0f584d^100^ with their in-house database made with MERC, MMETSP and Uniclust50 (http://wwwuser.gwdg.de/~compbiol/metaeuk/). The annotation was then subsequently done with eggNOG-mapper v2.1.0^101^. The completeness and contamination of the MAGs were assessed with Busco v5.0.0^102^ and the eukaryotic lineage (255 genes). We determined their taxonomy by comparing the results of the EUKulele v1.0.3^89^ and EukCC v0.3^103^ along with homology comparisons with publicly available genomes not included in the previous tools by protein BLAST v2.10.0^104^.

### Co-occurrence interaction networks

Co-occurrence networks between the pro- and eukaryotic MAGs were constructed using an average of the distance matrices created from SparCC^105^, Spearman’s correlation and SpiecEasi^106^ where the networks were constructed using the ‘Meinshausen and Bühlmann (mb)’ method. Nodes with fewer than two degrees were discarded to identify cliques with three or more interactions, while negative edges were removed to visualize only mutualistic relationships. The matrix was visualised using the *igraph^107^* R package. The largest component from the overall co-occurrence network was determined using the *components* module of the *igraph* package. Null model hypothesis was tested by assessing the distribution of the node degree and the respective probabilities of the occurrence network against those simulating the Erdos-Renyi, Barabasi-Albert, Stochastic-block null models^108^.

### Phylogenomics and pangenomes

For the pangenome analyses, we collected all the bins taxonomically identified as *Polaromonas* spp. and used the pangenome workflow described by Meren *et al*. (http://merenlab.org/2016/11/08/pangenomics-v2/) using anvi’o^93^, along with NCBI^109^ refseq genomes for comparison and an outgroup from the closely related *Rhodoferax* genus. The choice of *Polaromonas* spp. was based on its high abundance and prevalence within the epilithic biofilms. The accession IDs from the reference genomes obtained from NCBI are provided in the supplementary material. The pangenome was run using the *--min-bit 0.5*, *--mcl-inflation 10* and *--min-occurence 2* parameters, excluding the partial gene calls. A phylogenomic tree was built using MUSCLE v3.8.1551^110^ and FastTree2 v2.1.10^111^ on all single-copy gene clusters in the pangenome that were present in at least 30 genomes and had a functional homogeneity index below 0.9, and geometric homogeneity index above 0.9. The phylogenomic tree was used to order the genomes, the frequency of gene clusters (GC) to order the GC dendrogram. A phylogenomic bacterial tree of life containing the 47 high-quality MAGs along with 264 NCBI bacterial genomes was built based on a set of 74 single-copy genes using the GToTree v1.5.51^112^ pipeline with the -D parameter, allowing to retrieve taxonomic information for the NCBI accessions. Briefly, HMMER3 v3.3.2^113^ was used to retrieve the single-copy genes after gene-calling with Prodigal v2.6.3^114^ and aligned using TrimAl v1.4.rev15^115^. The entire workflow is based on GNU Parallel v20210222^116^.

### Data analyses and figures

Figures for the study including visualizations derived from the taxonomic and functional components, were created using version 3.6 of the R statistical software package^117^. The maps indicating the collection sites were generated using the *ggmap^118^* package in R. KEGGDecoder^119^ was used to assess enriched KEGG orthology (KO) IDs in comparison to 105 publicly available metagenome sampled in various ecosystems at a global scale (Supp. Tables 3 and 6), which were processed using the IMP workflow. *DESeq2^120^* with FDR-adjustments for multiple testing were used to assess KOs significantly enriched in the GFS metagenomes compared to this comparison dataset. The volcano plot highlighting the significant KOs was generated using the *EnhancedVolcano*^121^ R package. Figures from metabarcoding data were also generated in Rv3.6 using the *ggplot2*^122^ package and were further annotated graphically using Inkscape.

## Data availability

Raw sequencing data samples and the MAGs are available at NCBI’s sequence read archive under BioProject accession **PRJNA733707**. The Biosample accession IDs and the metadata associated with each sample are listed under Supp. Table 5.

## Competing interests

The authors declare no conflicts of interest.

## Code availability

The detailed code used for the for the downstream functional and growth analyses is available at https://git-r3lab.uni.lu/susheel.busi/nomis_pipeline. The custom pipeline for eukaryote analyses can be found here: https://github.com/Mass23/NOMIS_ENSEMBLE/tree/coassembly. Subsequent binning and manual refinement of eukaryotic MAGs was done as described here: https://git-r3lab.uni.lu/susheel.busi/nomis_pipeline/-/blob/master/workflow/notes/MiscEUKMAGs.md A snippet of the relevant results have been uploaded to Zenodo at https://doi.org/10.5281/zenodo.5545722.

## Funding

This research was funded by The NOMIS Foundation to TJB. SBB was supported by the Synergia grant (CRSII5_180241: Swiss National Science Foundation) to TJB. LdN and PW are supported by the Luxembourg National Research Fund (FNR; PRIDE17/11823097). RM and DD are supported by King Abdullah University of Science and Technology through the baseline research funds to DD.

## Acknowledgements

We are thankful for the assistance of Audrey Frachet Bour, Lea Grandmougin, Janine Habier, Laura Lebrun (LCSB) and Emmy Marie Oppliger (EPFL) for laboratory support. We are grateful to Alex Washburne for his feedback on the draft, and we also acknowledge the valuable input from Rashi Halder at the LCSB Sequencing Platform with respect to library preparation. We are equally grateful for the valuable insights into metagenomic processing from Patrick May and Cedric Christian Laczny, and especially Valentina Galata with the python scripts and Snakemake workflows. The computational analyses presented in this paper were carried out using the HPC facilities at the University of Luxembourg (https://hpc.uni.lu)^123^.

## Author contributions

SBB, MB, SF, HP, PW, and TJB conceived of the project. MiST, MT, VDS, MaSc, and HP conducted the fieldwork. PP and EMO extracted DNA, SBB and PP prepared the metagenomic and metabarcoding libraries, and RM and DD performed the sequencing. SBB conceptualized the data analyses, while SBB, MB, SF, GM performed the analyses. LdN contributed to the python scripts and Snakemake workflows for the analyses. SBB, MB, TJK, PW and TJB wrote the manuscript with significant input and editing from all coauthors.

---

**Supplementary Figure 1.**
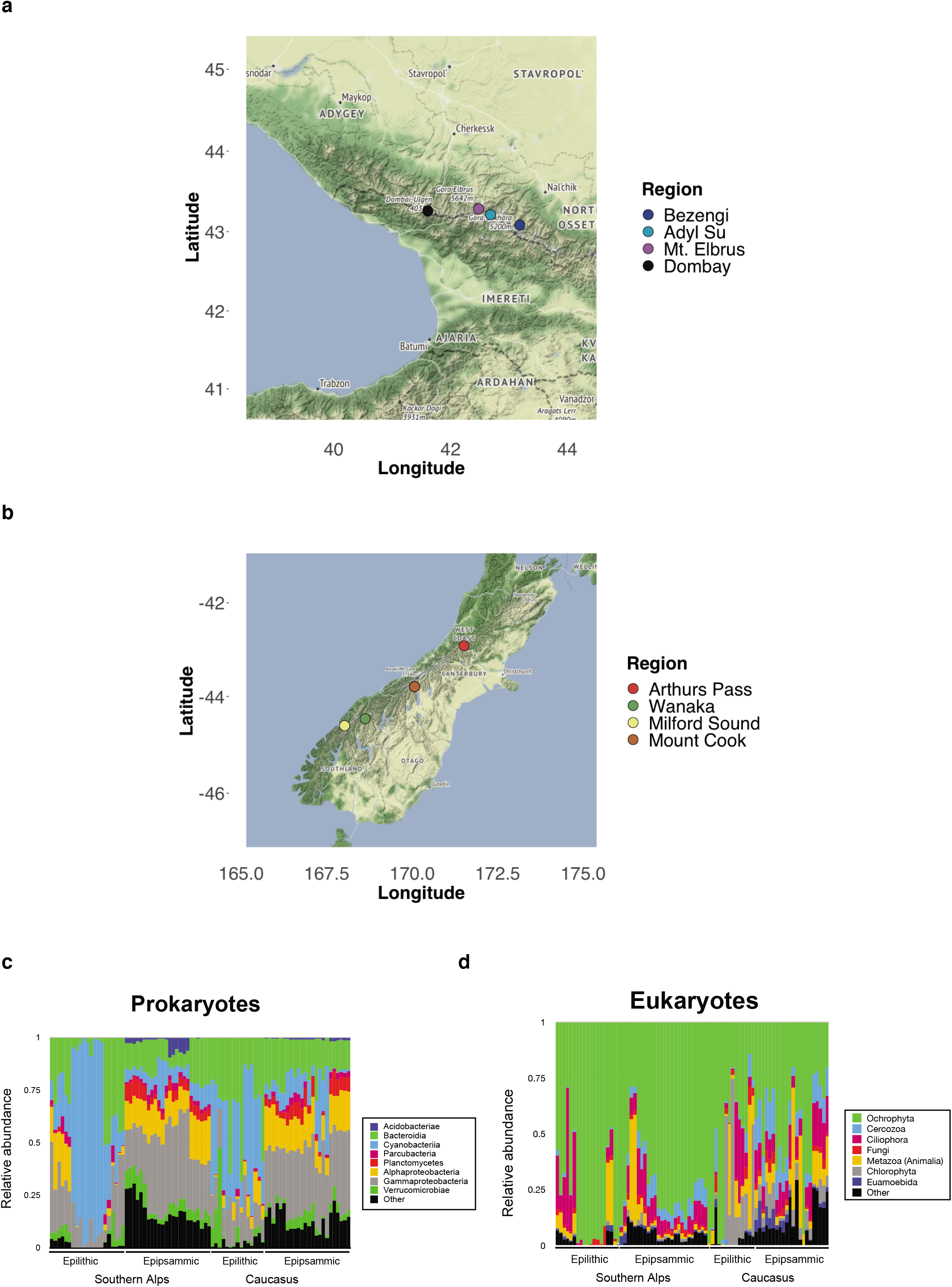
Sediment and epilithic biofilm sites. Regions indicating the collection sites for the epilithic and epipsammic biofilms from (a) Caucasus and (b) Southern Alps. Relative abundance of prokaryotes (c) and eukaryotes (d) at the phylum and subdomain levels based on the sequencing of the 16S and 18S rRNA genes, respectively.

**Supplementary Figure 2.**
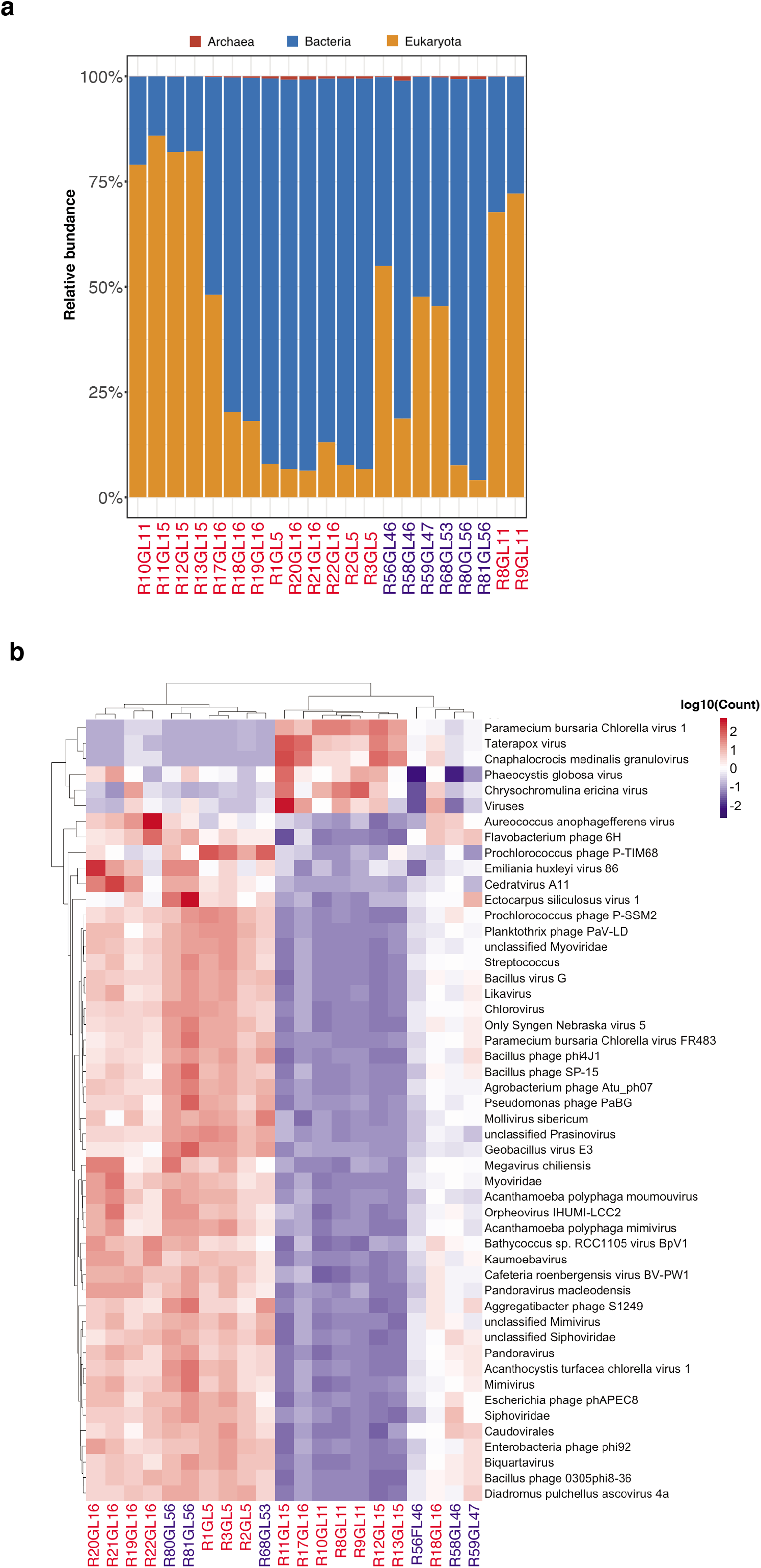
Epilithic biofilm metagenomic profiles. (a) Relative abundance profiles across the three domains of life: archaea, bacteria and eukaryotes in the epilithic biofilms, obtained from the sample metagenomes. Samples from the Southern Alps are indicated in red, while those from Caucasus are shown in blue. (b) Virome profile indicating the top 50 viruses. Scaled abundance from low (−2) to high (2) are indicated in the heatmap.

**Supplementary Figure 3.**
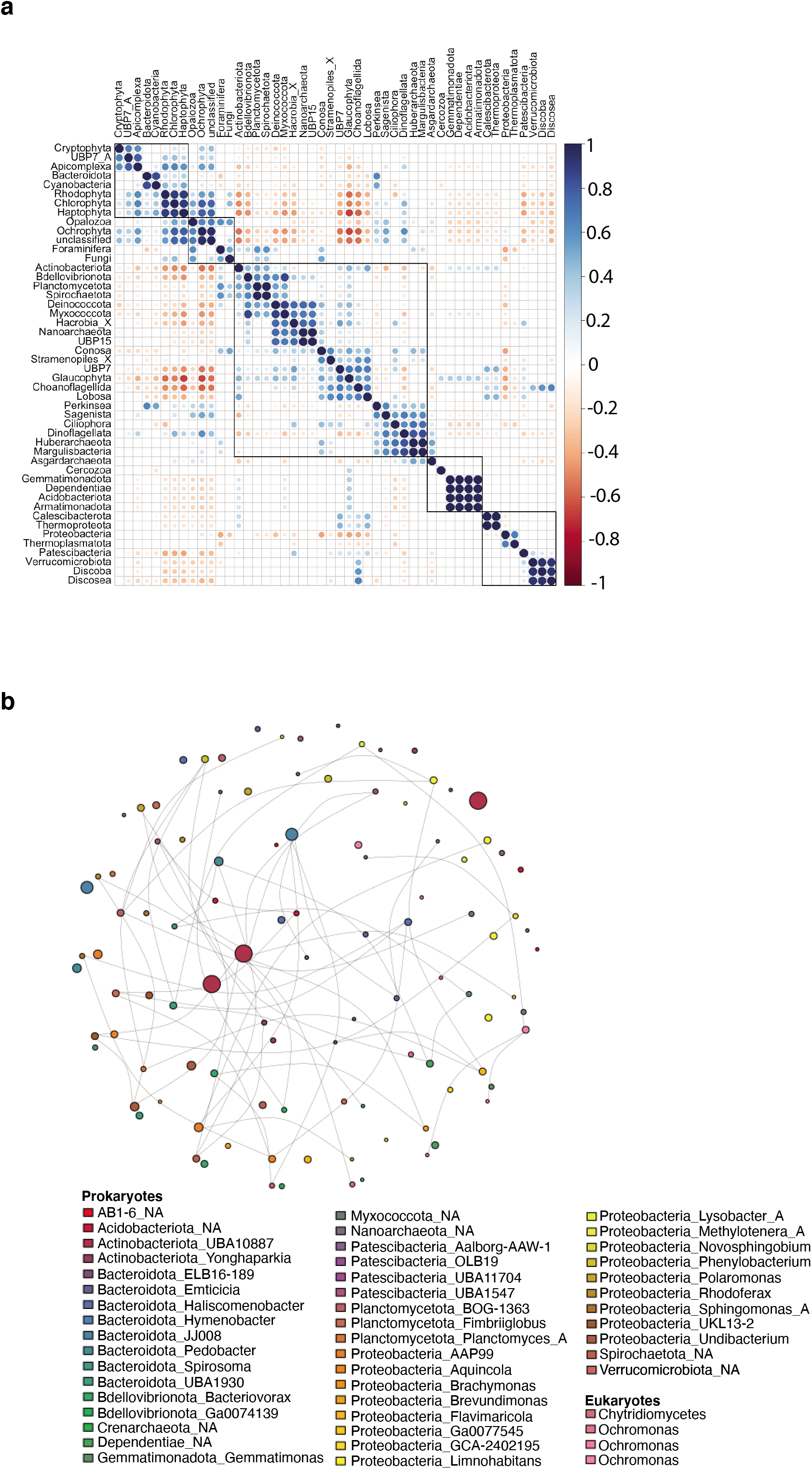
Cross-domain interactions and adaptations of epilithic biofilms. (a) Corrplot based on Spearman’s correlation between pro- and eukaryotic MAGs aggregated at the phylum level. (b) Co-occurrence network of all MAGs. Each node represents a MAG, while the size represents the degree centrality. The edges represent the positive coefficient of co-occurrence along with the corresponding betweenness centrality between the MAGs. Unconnected nodes represent MAGs with lower betweenness (< 0.5) compared to other MAGs. The color of the nodes represents the individual taxa, while the lines represent the edges connecting the nodes. The thickness of the lines indicates those edges with a betweenness greater than 0.5.

**Supplementary Figure 4.**
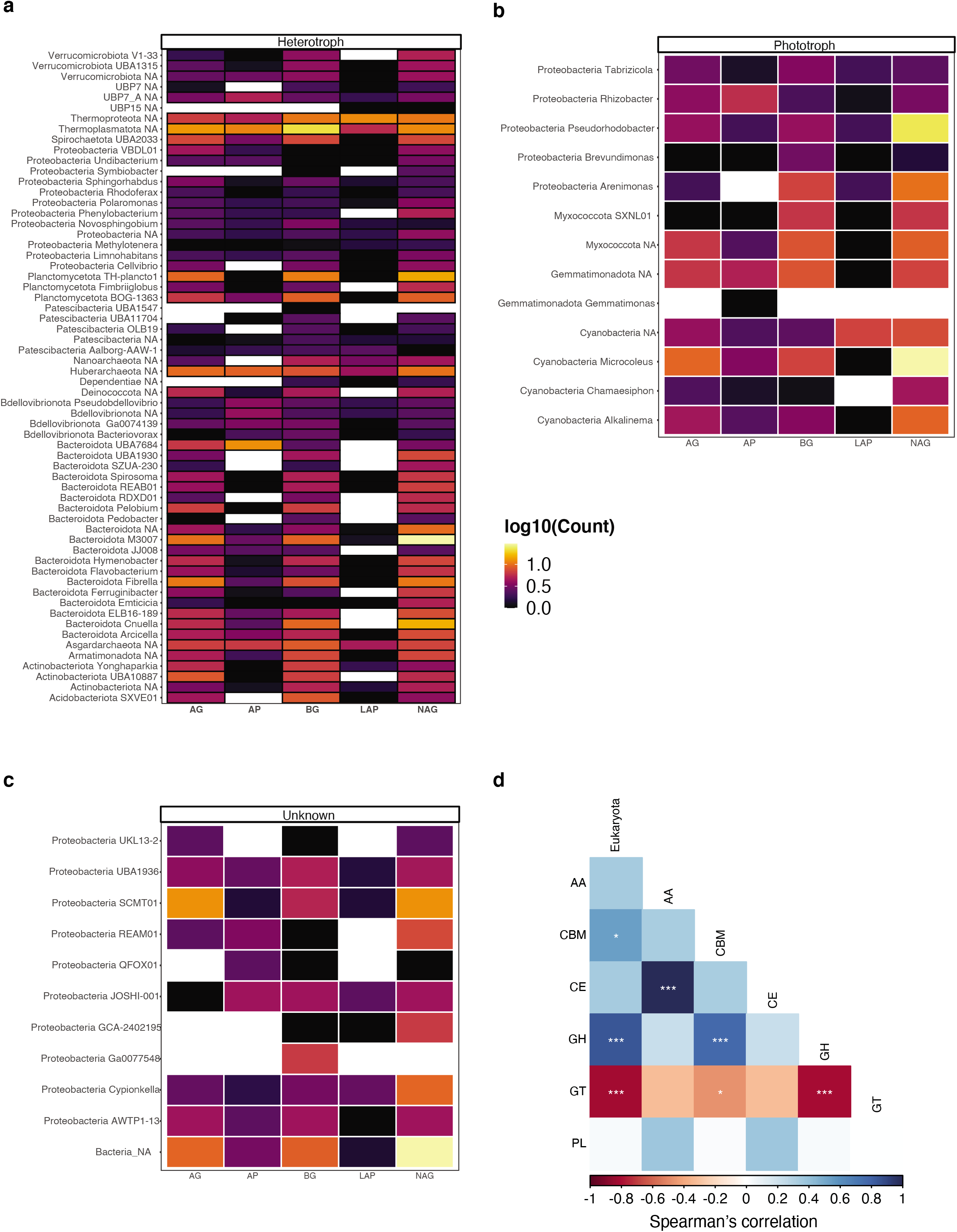
Extracellular enzyme genes based on lifestyle. The classification at phylum and genus levels of MAGs identified as (a) heterotrophs, (b) phototrophs, or (c) those with ‘unknown’ trophic metabolisms are depicted, showing the abundance of genes encoding for extracellular enzymes. NA: unclassified genus; AG: α-1,4-glucosidase; BG: β-1,4-glucosidase; LAP: leucine aminopeptidase; NAG: β-1,4-N-acetylglucosaminidase; AP: acid (alkaline) phosphatase. (d) (c) Spearman’s correlation analyses of overall eukaryote relative abundances with the CAZyme abundances. FDR-adjusted *p*-values are indicated by *, *i.e.*, * < 0.05, ** < 0.01, *** < 0.001.

**Supplementary Figure 5.**
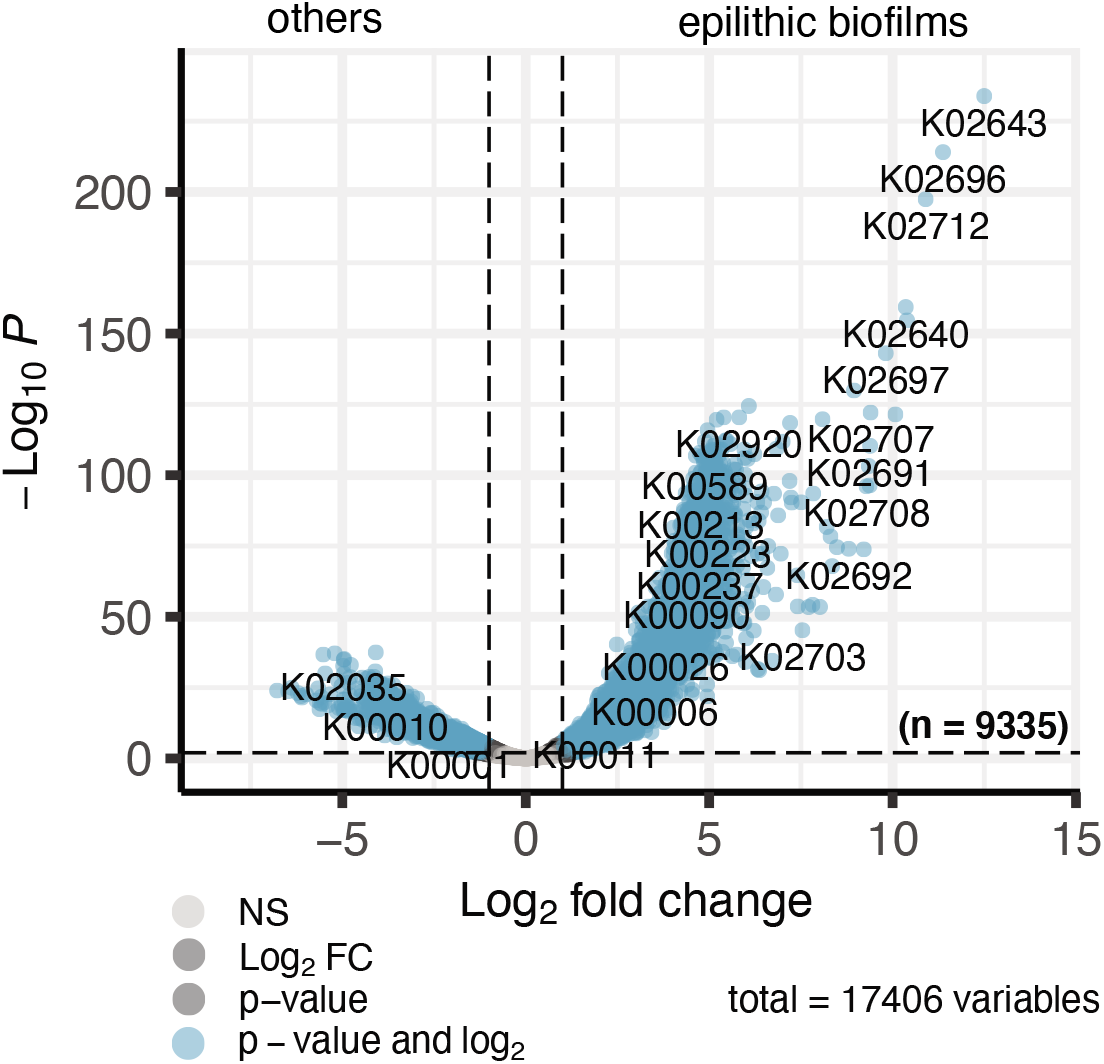
Comparison to public metagenomes reveals differential gene abundances. Volcano plot indicating the total number of KOs (n = 9,335; total = 17,406) enriched in epilithic biofilms compared to 105 publicly available metagenomes.

## Notes

https://doi.org/10.5281/zenodo.5545722

### Competing Interest Statement

The authors have declared no competing interest.

## References

1. Uehlinger, U., Robinson, C. T., Hieber, M. & Zah, R. The physico-chemical habitat template for periphyton in alpine glacial streams under a changing climate. in Global Change and River Ecosystems—Implications for Structure, Function and EcosystemServices (eds. Stevenson, R. J. & Sabater, S.) 107–121 (Springer Netherlands, 2010).

2. Battin, T. J., Wille, A., Psenner, R. & Richter, A. Large-scale environmental controls on microbial biofilms in high-alpine streams. Biogeosciences 1, 159–171 (2004).

3. Boix Canadell, M. et al. Regimes of primary production and their drivers in Alpine streams. Freshw. Biol. 66, 1449–1463 (2021).

4. Bernhardt, E. S. et al. Control Points in Ecosystems: Moving Beyond the Hot Spot Hot Moment Concept. Ecosystems 20, 665–682 (2017).

5. Huss, M. & Hock, R. Global-scale hydrological response to future glacier mass loss. Nat. Clim. Chang. 8, 135–140 (2018).

6. Milner, A. M. et al. Glacier shrinkage driving global changes in downstream systems. Proc. Natl. Acad. Sci. U. S. A. 114, 9770–9778 (2017).

7. Roncoroni, M., Brandani, J., Battin, T. I. & Lane, S. N. Ecosystem engineers: Biofilms and the ontogeny of glacier floodplain ecosystems. WIREs Water 6, e1390 (2019).

8. Hoyle, J. T., Kilroy, C., Hicks, D. M. & Brown, L. The influence of sediment mobility and channel geomorphology on periphyton abundance. Freshw. Biol. 62, 258–273 (2017).

9. Cole, J. J. Interactions Between Bacteria and Algae in Aquatic Ecosystems. Annu. Rev. Ecol. Syst. 13, 291–314 (1982).

10. Seymour, J. R., Amin, S. A., Raina, J.-B. & Stocker, R. Zooming in on the phycosphere: the ecological interface for phytoplankton–bacteria relationships. Nature Microbiology vol. 2 (2017).

11. Amin, S. A. et al. Interaction and signalling between a cosmopolitan phytoplankton and associated bacteria. Nature 522, 98–101 (2015).

12. Christie-Oleza, J. A., Sousoni, D., Lloyd, M., Armengaud, J. & Scanlan, D. J. Nutrient recycling facilitates long-term stability of marine microbial phototroph–heterotroph interactions. Nature Microbiology 2, 1–10 (2017).

13. Haack, T. K. & McFeters, G. A. Nutritional relationships among microorganisms in an epilithic biofilm community. Microb. Ecol. 8, 115–126 (1982).

14. Kaplan, L. A. & Bott, T. L. Diel fluctuations in bacterial activity on streambed substrata during vernal algal blooms: Effects of temperature, water chemistry, and habitat. Limnol. Oceanogr. 34, 718–733 (1989).

15. Vincent, W. F., Downes, M. T., Castenholz, R. W. & Howard-Williams, C. Community structure and pigment organisation of cyanobacteria-dominated microbial mats in Antarctica. European Journal of Phycology vol. 28 213–221 (1993).

16. Besemer, K., Singer, G., Hödl, I. & Battin, T. J. Bacterial community composition of stream biofilms in spatially variable-flow environments. Appl. Environ. Microbiol. 75, 7189–7195 (2009).

17. Risse‐Buhl, U. et al. Near streambed flow shapes microbial guilds within and across trophic levels in fluvial biofilms. Limnol. Oceanogr. 65, 2261–2277 (2020).

18. Palmer, M. A., Swan, C. M., Nelson, K., Silver, P. & Alvestad, R. Streambed landscapes: evidence that stream invertebrates respond to the type and spatial arrangement of patches. Landsc. Ecol. 15, 563–576 (2000).

19. Battin, T. J. et al. Microbial landscapes: new paths to biofilm research. Nat. Rev. Microbiol. 5, 76–81 (2007).

20. Dzubakova, K. et al. Environmental heterogeneity promotes spatial resilience of phototrophic biofilms in streambeds. Biol. Lett. 14, (2018).

21. Sloan, W. T. et al. Quantifying the roles of immigration and chance in shaping prokaryote community structure. Environ. Microbiol. 8, 732–740 (2006).

22. Hug, L. A. et al. A new view of the tree of life. Nat Microbiol 1, 16048 (2016).

23. Chaudhari, N. M., Overholt, W. A. & Figueroa-Gonzalez, P. A. The economical lifestyle of CPR bacteria in groundwater allows little preference for environmental drivers. bioRxiv (2021).

24. Vigneron, A. et al. Ultra‐small and abundant: Candidate phyla radiation bacteria are potential catalysts of carbon transformation in a thermokarst lake ecosystem. Limnol. Oceanogr. Lett. 5, 212–220 (2020).

25. Liu, Y. et al. Expanded diversity of Asgard archaea and their relationships with eukaryotes. Nature 593, 553–557 (2021).

26. Cai, M. et al. Ecological features and global distribution of Asgard archaea. Sci. Total Environ. 758, 143581 (2021).

27. Niedrist, G. H. & Füreder, L. When the going gets tough, the tough get going: The enigma of survival strategies in harsh glacial stream environments. Freshwater Biology vol. 63 1260–1272 (2018).

28. Bengtsson, M. M., Wagner, K., Schwab, C., Urich, T. & Battin, T. J. Light availability impacts structure and function of phototrophic stream biofilms across domains and trophic levels. Mol. Ecol. 27, 2913–2925 (2018).

29. Payne, A. T. et al. Widespread cryptic viral infections in lotic biofilms. Biofilms 2, 100016 (2020).

30. Anesio, A. M., Mindl, B., Laybourn-Parry, J., Hodson, A. J. & Sattler, B. Viral dynamics in cryoconite holes on a high Arctic glacier (Svalbard). J. Geophys. Res. 112, (2007).

31. Bellas, C. M., Schroeder, D. C., Edwards, A., Barker, G. & Anesio, A. M. Flexible genes establish widespread bacteriophage pan-genomes in cryoconite hole ecosystems. Nat. Commun. 11, 4403 (2020).

32. Liu, Q. et al. Light stimulates anoxic and oligotrophic growth of glacial Flavobacterium strains that produce zeaxanthin. ISME J. 15, 1844–1857 (2021).

33. Sánchez Barranco, V. et al. Trophic position, elemental ratios and nitrogen transfer in a planktonic host-parasite-consumer food chain including a fungal parasite. Oecologia 194, 541–554 (2020).

34. Klawonn, I. et al. Characterizing the ‘fungal shunt’: Parasitic fungi on diatoms affect carbon flow and bacterial communities in aquatic microbial food webs. Proc. Natl. Acad. Sci. U. S. A. 118, (2021).

35. Sinsabaugh, R. L., Hill, B. H. & Follstad Shah, J. J. Ecoenzymatic stoichiometry of microbial organic nutrient acquisition in soil and sediment. Nature 462, 795–798 (2009).

36. Avcı, B., Krüger, K., Fuchs, B. M., Teeling, H. & Amann, R. I. Polysaccharide niche partitioning of distinct Polaribacter clades during North Sea spring algal blooms. ISME J. 14, 1369–1383 (2020).

37. Sichert, A. et al. Verrucomicrobia use hundreds of enzymes to digest the algal polysaccharide fucoidan. Nat Microbiol 5, 1026–1039 (2020).

38. Zhou, J., Lyu, Y., Richlen, M., Anderson, D. M. & Cai, Z. Quorum sensing is a language of chemical signals and plays an ecological role in algal-bacterial interactions. CRC Crit. Rev. Plant Sci. 35, 81–105 (2016).

39. Croft, M. T., Lawrence, A. D., Raux-Deery, E., Warren, M. J. & Smith, A. G. Algae acquire vitamin B12 through a symbiotic relationship with bacteria. Nature 438, 90–93 (2005).

40. Grossman, A. Nutrient Acquisition: The Generation of Bioactive Vitamin B12 by Microalgae. Current biology: CB vol. 26 R319–21 (2016).

41. Segev, E. et al. Dynamic metabolic exchange governs a marine algal-bacterial interaction. Elife 5, (2016).

42. Anesio, A. M., Lutz, S., Chrismas, N. A. M. & Benning, L. G. The microbiome of glaciers and ice sheets. NPJ Biofilms Microbiomes 3, 10 (2017).

43. Tranter, M., Mills, R. & Raiswell, R. Chemical weathering reactions in Alpine glacial meltwaters. in International symposium on water-rock interaction 687–690 (1989).

44. Tranter, M., Brown, G., Raiswell, R., Sharp, M. & Gurnell, A. A conceptual model of solute acquisition by Alpine glacial meltwaters. J. Glaciol. 39, 573–581 (1993).

45. St Pierre, K. A. et al. Proglacial freshwaters are significant and previously unrecognized sinks of atmospheric CO2. Proc. Natl. Acad. Sci. U. S. A. 116, 17690–17695 (2019).

46. Dunham, E. C., Dore, J. E., Skidmore, M. L., Roden, E. E. & Boyd, E. S. Lithogenic hydrogen supports microbial primary production in subglacial and proglacial environments. Proc. Natl. Acad. Sci. U. S. A. 118, (2021).

47. Hernández, M. et al. Reconstructing Genomes of Carbon Monoxide Oxidisers in Volcanic Deposits Including Members of the Class Ktedonobacteria. Microorganisms 8, (2020).

48. Quick, A. M. et al. Nitrous oxide from streams and rivers: A review of primary biogeochemical pathways and environmental variables. Earth-Sci. Rev. 191, 224–262 (2019).

49. Kuypers, M. M. M., Marchant, H. K. & Kartal, B. The microbial nitrogen-cycling network. Nat. Rev. Microbiol. 16, 263–276 (2018).

50. Gooseff, M. N., McKnight, D. M., Runkel, R. L. & Duff, J. H. Denitrification and hydrologic transient storage in a glacial meltwater stream, McMurdo Dry Valleys, Antarctica. Limnol. Oceanogr. 49, 1884–1895 (2004).

51. Varin, T., Lovejoy, C., Jungblut, A. D., Vincent, W. F. & Corbeil, J. Metagenomic profiling of Arctic microbial mat communities as nutrient scavenging and recycling systems. Limnol. Oceanogr. 55, 1901–1911 (2010).

52. Kohler, T. J. et al. Patterns and Drivers of Extracellular Enzyme Activity in New Zealand Glacier-Fed Streams. Front. Microbiol. 11, 591465 (2020).

53. Alves, R. J. E. et al. Ammonia Oxidation by the Arctic Terrestrial Thaumarchaeote Candidatus Nitrosocosmicus arcticus Is Stimulated by Increasing Temperatures. Front. Microbiol. 10, 1571 (2019).

54. Könneke, M. et al. Ammonia-oxidizing archaea use the most energy-efficient aerobic pathway for CO2 fixation. Proc. Natl. Acad. Sci. U. S. A. 111, 8239–8244 (2014).

55. Cockell, C. S. et al. Influence of ice and snow covers on the UV exposure of terrestrial microbial communities: dosimetric studies. J. Photochem. Photobiol. B 68, 23–32 (2002).

56. Sommaruga, R. The role of solar UV radiation in the ecology of alpine lakes. J. Photochem. Photobiol. B 62, 35–42 (2001).

57. Margesin, R. & Collins, T. Microbial ecology of the cryosphere (glacial and permafrost habitats): current knowledge. Appl. Microbiol. Biotechnol. 103, 2537–2549 (2019).

58. De Maayer, P., Anderson, D., Cary, C. & Cowan, D. A. Some like it cold: understanding the survival strategies of psychrophiles. EMBO Rep. 15, 508–517 (2014).

59. Tribelli, P. M. & López, N. I. Reporting Key Features in Cold-Adapted Bacteria. Life 8, (2018).

60. Varin, T., Lovejoy, C., Jungblut, A. D., Vincent, W. F. & Corbeil, J. Metagenomic analysis of stress genes in microbial mat communities from Antarctica and the High Arctic. Appl. Environ. Microbiol. 78, 549–559 (2012).

61. Alonso-Sáez, L. et al. Winter bloom of a rare betaproteobacterium in the Arctic Ocean. Front. Microbiol. 5, 425 (2014).

62. Hornung, C. et al. The Janthinobacterium sp. HH01 genome encodes a homologue of the V. cholerae CqsA and L. pneumophila LqsA autoinducer synthases. PLoS One 8, e55045 (2013).

63. Maillot, N. J., Honoré, F. A., Byrne, D., Méjean, V. & Genest, O. Cold adaptation in the environmental bacterium Shewanella oneidensis is controlled by a J-domain co-chaperone protein network. Commun Biol 2, 323 (2019).

64. Konings, W. N., Albers, S.-V., Koning, S. & Driessen, A. J. M. The cell membrane plays a crucial role in survival of bacteria and archaea in extreme environments. Antonie Van Leeuwenhoek 81, 61–72 (2002).

65. Methé, B. A. et al. The psychrophilic lifestyle as revealed by the genome sequence of Colwellia psychrerythraea 34H through genomic and proteomic analyses. Proc. Natl. Acad. Sci. U. S. A. 102, 10913–10918 (2005).

66. Metagenomic Analysis of Stress Genes in Microbial Mat Communities from Antarctica and the High Arctic. https://journals.asm.org/doi/abs/10.1128/AEM.06354-11.

67. Ayala-del-Río, H. L. et al. The genome sequence of Psychrobacter arcticus 273-4, a psychroactive Siberian permafrost bacterium, reveals mechanisms for adaptation to low-temperature growth. Appl. Environ. Microbiol. 76, 2304–2312 (2010).

68. Mykytczuk, N. C. S. et al. Bacterial growth at −15 °C; molecular insights from the permafrost bacterium Planococcus halocryophilus Or1. The ISME Journal vol. 7 1211–1226 (2013).

69. Busi, S. B. et al. Optimised biomolecular extraction for metagenomic analysis of microbial biofilms from high-mountain streams. PeerJ 8, e9973 (2020).

70. Fodelianakis, S. et al. Microdiversity characterizes prevalent phylogenetic clades in the glacier-fed stream microbiome. ISME J. (2021) doi:10.1038/s41396-021-01106-6.

71. Stoeck, T. et al. Multiple marker parallel tag environmental DNA sequencing reveals a highly complex eukaryotic community in marine anoxic water. Mol. Ecol. 19 Suppl 1, 21–31 (2010).

72. Bolger, A. M., Lohse, M. & Usadel, B. Trimmomatic: a flexible trimmer for Illumina sequence data. Bioinformatics 30, 2114–2120 (2014).

73. Bolyen, E. et al. Reproducible, interactive, scalable and extensible microbiome data science using QIIME 2. Nat. Biotechnol. 37, 852–857 (2019).

74. Quast, C. et al. The SILVA ribosomal RNA gene database project: improved data processing and web-based tools. Nucleic Acids Res. 41, D590–6 (2013).

75. Dixon, P. VEGAN, a package of R functions for community ecology. Journal of Vegetation Science vol. 14 927–930 (2003).

76. Foster, Z. S. L., Sharpton, T. J. & Grünwald, N. J. Metacoder: An R package for visualization and manipulation of community taxonomic diversity data. PLoS Comput. Biol. 13, e1005404 (2017).

77. Burns, A. R. et al. Contribution of neutral processes to the assembly of gut microbial communities in the zebrafish over host development. ISME J. 10, 655–664 (2016).

78. Narayanasamy, S. et al. IMP: a pipeline for reproducible reference-independent integrated metagenomic and metatranscriptomic analyses. Genome Biol. 17, 260 (2016).

79. Heintz-Buschart, A. et al. Integrated multi-omics of the human gut microbiome in a case study of familial type 1 diabetes. Nat Microbiol 2, 16180 (2016).

80. Li, D., Liu, C.-M., Luo, R., Sadakane, K. & Lam, T.-W. MEGAHIT: an ultra-fast single-node solution for large and complex metagenomics assembly via succinct de Bruijn graph. Bioinformatics 31, 1674–1676 (2015).

81. Kang, D. D. et al. MetaBAT 2: an adaptive binning algorithm for robust and efficient genome reconstruction from metagenome assemblies. PeerJ 7, e7359 (2019).

82. Wu, Y.-W., Simmons, B. A. & Singer, S. W. MaxBin 2.0: an automated binning algorithm to recover genomes from multiple metagenomic datasets. Bioinformatics 32, 605–607 (2016).

83. Sieber, C. M. K. et al. Recovery of genomes from metagenomes via a dereplication, aggregation and scoring strategy. Nat Microbiol 3, 836–843 (2018).

84. Parks, D. H., Imelfort, M., Skennerton, C. T., Hugenholtz, P. & Tyson, G. W. CheckM: assessing the quality of microbial genomes recovered from isolates, single cells, and metagenomes. Genome Res. 25, 1043–1055 (2015).

85. Chaumeil, P.-A., Mussig, A. J., Hugenholtz, P. & Parks, D. H. GTDB-Tk: a toolkit to classify genomes with the Genome Taxonomy Database. Bioinformatics (2019) doi:10.1093/bioinformatics/btz848.

86. Kieft, K., Zhou, Z. & Anantharaman, K. VIBRANT: automated recovery, annotation and curation of microbial viruses, and evaluation of viral community function from genomic sequences. Microbiome 8, 90 (2020).

87. Zablocki, O., Jang, H. B., Bolduc, B. & Sullivan, M. B. vConTACT 2: A Tool to Automate Genome-Based Prokaryotic Viral Taxonomy. in Plant and Animal Genome XXVII Conference (January 12–16, 2019) (PAG, 2019).

88. Nayfach, S. et al. CheckV assesses the quality and completeness of metagenome-assembled viral genomes. Nat. Biotechnol. 39, 578–585 (2021).

89. Krinos, A. I., Hu, S. K., Cohen, N. R. & Alexander, H. EUKulele: Taxonomic annotation of the unsung eukaryotic microbes. arXiv [q-bio.PE] (2020).

90. Queirós, P., Delogu, F., Hickl, O., May, P. & Wilmes, P. Mantis: flexible and consensus-driven genome annotation. bioRxiv (2020).

91. Zhou, Z. et al. METABOLIC: High-throughput profiling of microbial genomes for functional traits, biogeochemistry, and community-scale metabolic networks. bioRxiv 761643 (2020) doi:10.1101/761643.

92. McDaniel, E. A., Anantharaman, K. & McMahon, K. D. metabolisHMM: Phylogenomic analysis for exploration of microbial phylogenies and metabolic pathways. bioRxiv 2019.12.20.884627 (2019) doi:10.1101/2019.12.20.884627.

93. Eren, A. M. et al. Anvi’o: an advanced analysis and visualization platform for ‘omics data. PeerJ 3, e1319 (2015).

94. Weissman, J. L., Hou, S. & Fuhrman, J. A. Estimating maximal microbial growth rates from cultures, metagenomes, and single cells via codon usage patterns. Proc. Natl. Acad. Sci. U. S. A. 118, (2021).

95. Chen, S., Zhou, Y., Chen, Y. & Gu, J. fastp: an ultra-fast all-in-one FASTQ preprocessor. Bioinformatics 34, i884–i890 (2018).

96. Bushnell, B. BBMap: A fast, accurate, splice-aware aligner. https://www.osti.gov/biblio/1241166 (2014).

97. Wood, D. E., Lu, J. & Langmead, B. Improved metagenomic analysis with Kraken 2. Genome Biol. 20, 257 (2019).

98. West, P. T., Probst, A. J., Grigoriev, I. V., Thomas, B. C. & Banfield, J. F. Genome-reconstruction for eukaryotes from complex natural microbial communities. Genome Res. 28, 569–580 (04 2018).

99. Alneberg, J. et al. CONCOCT: Clustering cONtigs on COverage and ComposiTion. arXiv [q-bio.GN] (2013).

100. Levy Karin, E., Mirdita, M. & Söding, J. MetaEuk-sensitive, high-throughput gene discovery, and annotation for large-scale eukaryotic metagenomics. Microbiome 8, 48 (2020).

101. Huerta-Cepas, J. et al. eggNOG 5.0: a hierarchical, functionally and phylogenetically annotated orthology resource based on 5090 organisms and 2502 viruses. Nucleic Acids Res. 47, D309–D314 (2019).

102. Seppey, M., Manni, M. & Zdobnov, E. M. BUSCO: Assessing Genome Assembly and Annotation Completeness. Methods Mol. Biol. 1962, 227–245 (2019).

103. Saary, P., Mitchell, A. L. & Finn, R. D. Estimating the quality of eukaryotic genomes recovered from metagenomic analysis with EukCC. Genome Biol. 21, 244 (2020).

104. Altschul, S. F., Gish, W., Miller, W., Myers, E. W. & Lipman, D. J. Basic local alignment search tool. J. Mol. Biol. 215, 403–410 (1990).

105. Inferring Correlation Networks from Genomic Survey Data. https://journals.plos.org/ploscompbiol/article?id=10.1371/journal.pcbi.1002687.

106. Kurtz, Z. D. et al. Sparse and compositionally robust inference of microbial ecological networks. PLoS Comput. Biol. 11, e1004226 (2015).

107. Csardi, G. & Nepusz, T. The igraph software package for complex network research. InterJournal, Complex Systems 1695, 1–9 (2006).

108. Dormann, C. F., Frund, J., Bluthgen, N. & Gruber, B. Indices, graphs and null models: Analyzing bipartite ecological networks. Open Ecol. J. 2, 7–24 (2009).

109. Pruitt, K. D., Tatusova, T. & Maglott, D. R. NCBI reference sequences (RefSeq): a curated non-redundant sequence database of genomes, transcripts and proteins. Nucleic Acids Res. 35, D61–5 (2007).

110. Edgar, R. C. MUSCLE: multiple sequence alignment with high accuracy and high throughput. Nucleic Acids Res. 32, 1792–1797 (2004).

111. Price, M. N., Dehal, P. S. & Arkin, A. P. FastTree 2--approximately maximum-likelihood trees for large alignments. PLoS One 5, e9490 (2010).

112. Lee, M. D. GToTree: a user-friendly workflow for phylogenomics. Bioinformatics 35, 4162–4164 (2019).

113. Eddy, S. R. Accelerated Profile HMM Searches. PLoS Comput. Biol. 7, e1002195 (2011).

114. Hyatt, D. et al. Prodigal: prokaryotic gene recognition and translation initiation site identification. BMC Bioinformatics 11, 119 (2010).

115. Capella-Gutiérrez, S., Silla-Martínez, J. M. & Gabaldón, T. trimAl: a tool for automated alignment trimming in large-scale phylogenetic analyses. Bioinformatics 25, 1972–1973 (2009).

116. Tange, O. GNU Parallel 2018. (https://Lulu.com, 2018).

117. Team, R. C. & Others. R: A language and environment for statistical computing. (2013).

118. Kahle, D. & Wickham, H. Ggmap: Spatial visualization with ggplot2. R J. 5, 144 (2013).

119. Graham, E. D., Heidelberg, J. F. & Tully, B. J. Potential for primary productivity in a globally-distributed bacterial phototroph. ISME J. 12, 1861–1866 (2018).

120. Love, M. I., Huber, W. & Anders, S. Moderated estimation of fold change and dispersion for RNA-seq data with DESeq2. Genome Biol. 15, (2014).

121. kevinblighe/EnhancedVolcano: Publication-ready volcano plots with enhanced colouring and labeling. https://github.com/kevinblighe/EnhancedVolcano.

122. Wickham, H. ggplot2: ggplot2. Wiley Interdiscip. Rev. Comput. Stat. 3, 180–185 (2011).

123. Varrette, S., Bouvry, P., Cartiaux, H. & Georgatos, F. Management of an academic HPC cluster: The UL experience. in 2014 International Conference on High Performance Computing Simulation (HPCS) 959–967 (2014).

