## Supplementary text for "Genomic and metabolic adaptations of biofilms to ecological windows of opportunities in glacier-fed streams"

**Genomic insights into microbial islands of life in alpine streams**

Susheel Bhanu Busi^1^, Massimo Bourquin^2^, Stilianos Fodelianakis^2^, Grégoire Michoud^2^, Hannes Peter^2^, Tyler J. Kohler^2^, Paraskevi Pramateftaki^2^, Valentina Galata^1^, Michail Styllas^2^, Matteo Tolosano^2^, Vincent De Staercke^2^, Martina Schön^2^, Jade Brandani^2^, Leila Ezzat^2^, Emmy Marie Oppliger^2^, Alex D. Washburne^3,4^, Ramona Marasco^5^, Daniele Daffonchio^5^, Paul Wilmes^1,*^, & Tom J. Battin^2,*^

^1^Systems Ecology Group, Luxembourg Centre for Systems Biomedicine, University of Luxembourg, Esch-sur-Alzette, Luxembourg

^2^Stream Biofilm & Ecosystem Research Lab, ENAC Division, Ecole Polytechnique Federale de Lausanne, Lausanne, Switzerland

^3^Department of Microbiology and Immunology, Montana State University, USA

^4^Selva Analytics, LLC, Bozeman, Montana, USA

^5^Biological and Environmental Sciences and Engineering Division (BESE), King Abdullah University of Science and Technology (KAUST), Thuwal, Saudi Arabia

**A) DNA extraction protocol from alpine stream biofilms (rDNA)**

**Remark:** Every time you open the tubes make sure that there is no liquid on the lids by applying a short spin

1. In a 1.5-ml tube add 10-20% (~300 ul) 0.1 mm Zirconium beads (Cole-Parmer 36270-62) per volume and 750 ml of Lysis buffer mixed with 0.5ul of RNase (100 mg/ml, Qiagen 19101)
2. Add 0.05 to 0.1 g and bead-beat at 6000 r/min, 2x 15sec-break 15sec (Precellys 24 homogenizer)
3. Incubate at 37 °C for 1 h with gentle agitation
4. Spin samples, add 5 ul Proteinase K (20 mg/ml, Fisher Scientific Cat.No. 25530049) and mix a few times
5. Incubate statically at 70 °C for 10 min
6. Centrifuge at 12.000 x g for 1 min and transfer all supernatant to a new 1.5 ml microtube
7. Spin samples and add 1 vol of Phenol:CHCl3:IAA (Fisher Scientific, 15593049)
8. Mix thoroughly and centrifuge at 13.000 x g for 10 min
9. Transfer aqueous phase into a new 1.5-ml tube and add 1 vol ml Chloroform – isoamyl alcohol mixture (Sigma, 25666)
10. Mix thoroughly and centrifuge at 13.000 x g for 5 min
11. Transfer supernatant to a new 2ml tube and then add 1/10th volume of 3M sodium acetate (pH 5.2) (Sigma S7899)
12. Add 0.7 volumes of ice-cold Isopropanol (Sigma I9516) and mix thoroughly
13. Precipitate DNA at -20 °C overnight
14. Centrifuge at 12.000 x g at 4 °C for 15 min
15. Remove supernatant and discard without disturbing the pellet
16. Wash 2 times with 0.4 ml of 70% EtOH and centrifuge at 13.000 g at 4 °C for 10 min
17. Air-dry the pellet, and elute with 100 ul RNase-free, DNase-free water (Qiagen 129112)
18. Let DNA pellet to dissolve o/n at 4 °C
19. Use 2 ul sample to quantify DNA using Qubit HS dsDNA (Invitrogen Q32854)

**B) NCBI Accessions for *Polaromonas* genomes**

| **Name** | **AccessionID** |
| --- | --- |
| OUT1 | GCF_001955735.1_ASM195573v1 |
| DB1 | GCF_000013865.1_ASM1386v1 |
| DB2 | GCF_000015505.1_ASM1550v1 |
| DB3 | GCF_000282655.1_Polaromonas.strCF318_v1.0 |
| DB4 | GCF_000688115.1_ASM68811v1 |
| DB5 | GCF_000709345.1_Polaromonas_sp. |
| DB6 | GCF_001598235.1_ASM159823v1 |
| DB7 | GCF_002001015.1_ASM200101v1 |
| DB8 | GCF_002002705.1_ASM200270v1 |
| DB9 | GCF_002379085.1_ASM237908v1 |
| DB10 | GCF_002379095.1_ASM237909v1 |
| DB11 | GCF_003711205.1_ASM371120v1 |
| DB12 | GCF_009664225.1_ASM966422v1 |
| DB13 | GCF_012584515.1_ASM1258451v1 |
| DB14 | GCF_014641715.1_ASM1464171v1 |
| DB15 | GCF_015751795.1_ASM1575179v1 |
| DB16 | GCF_015752205.1_ASM1575220v1 |
| DB17 | GCF_015752225.1_ASM1575222v1 |
| DB18 | GCF_900103405.1_IMG-taxon_2636416056_annotated_assembly |
| DB19 | GCF_900112285.1_IMG-taxon_2609459740_annotated_assembly |
| DB20 | GCF_900116715.1_IMG-taxon_2615840640_annotated_assembly |

**C) Sloan model summary**

The dispersal rate coefficient (m) and the goodness of fit of the beta distribution model (R^2^) based on the Sloan neutral model analyses are indicated for New Zealand and Caucasus with respect to the metabarcoding information per amplicon sequence variant (ASV).

|  | 16S rRNA gene amplicons | | 18S rRNA gene amplicons | |
| --- | --- | --- | --- | --- |
|  | R^2^ | m | R^2^ | m |
| New Zealand | -0.726 | 3.7E-04±4E-04 | -0.204 | 4.3E-04±3E-04 |
| Caucasus | -1.11 | 6.4E-04±5E-04 | -0.527 | 0.57±0.31 |
